# *Shewanella* is a putative producer of polyunsaturated fatty acids in the gut soil of the composting earthworm *Eisenia fetida*

**DOI:** 10.1101/2024.03.31.587473

**Authors:** Jan-Philipp Wittlinger, Natalia Castejón, Bela Hausmann, David Berry, Stephanie L. Schnorr

## Abstract

Polyunsaturated fatty acids (PUFAs) play a crucial role in aiding bacteria to adapt to extreme and stressful environments. While there is a well-established understanding of their production, accrual, and transfer within marine ecosystems, knowledge about terrestrial environments remains limited. Investigation of the intestinal microbiome of earthworms has illuminated the presence of PUFAs presumably of microbial origin, which contrasts with the surrounding soil.

To comprehensively study this phenomenon, a multi-faceted approach was employed, combining fatty acid analysis with amplicon sequencing of the PfaA-KS domain of the anaerobic fatty acid synthase gene (*pfa*), as well as the 16S rRNA and 18S rRNA genes. This methodology was applied to scrutinize the gut microbiome of *Eisenia fetida*, its compost-based dietary source, and the resultant castings.

This study unveiled a distinct gut soil ecosystem from input compost and output castings in fatty acid profile as well as type and abundance of organisms. 16S sequencing provided insights into the microbial composition, showing increased relative abundance of certain Pseudomonadota, including *Shewanellaceae*, and Planctomycetota, including *Gemmataceae* within the gut microbiome compared to input bulk soil compost, while Actinomycetota and Bacillota were relatively enriched compared to the casted feces. Sequencing of the PfaA-KS domain revealed ASVs belonging primarily to *Shewanella*. Intriguingly, the 20C PUFAs were identified only in gut-soil samples, though PfaA-KS sequence abundance was highest in output castings, indicating a unique metabolism occurring only in the gut. Overall, the results indicate that *Shewanella* can explain PUFA enrichment in the gut environment because of *pfa* gene presence detected via PfaA-KS sequence data.

**Importance:** Prior research has demonstrated that earthworm microbiomes can potentially harbor PUFAs that are not found within their residing soil environment. Moreover, distinct indicator species have been pinpointed for various microbial genera in earthworm microbiomes. Nevertheless, none of these studies have integrated metataxonomic and fatty acid analysis to explore the origin of PUFA synthesis in any earthworm species, with the objective of identifying the specific organisms and locations responsible for this production. This study suggests that earthworms accumulate PUFAs produced from bacteria, especially *Shewanella,* activated through the gut ecosystem.

## Introduction

Fatty acids (FAs) are a key component of various lipid types and play important biological roles in life, such as serving as an energy resource, participating in signaling pathways, and stabilizing cell membranes (1–3). They are incorporated into lipid structures through ester bonds and can be classified into different groups based on their carbon chain length. Further they can be classified on their degree of saturation and the location and orientation of double bonds (4). Poly-unsaturated fatty acids (PUFAs), especially omega-3 and omega-6 PUFAs, are essential fatty acids for all higher organisms and have diverse functions, including building the lipid bilayer of plasma membranes (5–8). The degree of saturation of the lipid bilayer determines its permeability and fluidity, which can influence cell integrity and metabolism for energy production, growth and replication, particularly in response to environmental stressors like pressure, temperature, or oxidative stress (2, 9, 10). Omega-3 (ω3) fatty acids have anti-inflammatory functions and oppose the pro-inflammatory effects mediated by omega-6 (ω6) fatty acids. The omega-3 and omega-6 PUFAs, especially eicosapentaenoic acid (EPA), arachidonic acid (ARA), and docosahexaenoic acid (DHA), are physiologically important for broad precursors for eicosanoid secondary metabolites (8, 11). The production of eicosanoids is critical for insects and annelids to modulate physiological processes and immunity (12, 13).

### *De novo* PUFA synthesis by microorganisms

PUFAs are synthesized by two known pathways. The aerobic pathway involves elongation and oxygenic desaturation steps which are catalyzed by specific desaturases and elongases until the final PUFA length and unsaturation is reached (14). The anaerobic pathway, synthesizes PUFAs via a multifunctional mega-enzyme and double bonds are not introduced through oxygenic desaturation but rather by a PUFA synthase (15). This process, which was fully elucidated in 2001 (16), is encoded by a type I polyketide-like fatty acid synthase consisting of canonical fatty acid synthase domains (15), situated among either three genes (protozoa and myxobacteria) or five genes (marine bacteria). While the aerobic pathway is used by higher animals, lower plants, and eukaryotic microorganisms, the anaerobic pathway is limited to microorganisms such as fungi, algae, and bacteria (17).

The final fatty acid product that can be produced depends on the types of elongases and desaturases encoded in the genome and the selective conditions of their activity. A description of the enzymatic pathway and evolutionary considerations of organismal production patterns have been covered in detail, readers are especially referred to Kabeya et al. 2018 and Twining et al. 2021 (18, 19). To summarize, there is widespread presence of aerobic PUFA synthase genes among invertebrate taxa, including members of Nematoda, Cnidaria, Rotifera, Mollusca, Arthropoda, and Annelida. These genes putatively confer de novo synthesis capacity, and their functional characterization gives further evidence that their presence is part of an evolutionarily conserved adaptation to meet physiological needs for certain long-chain PUFAs. Even so, major groups of invertebrate producers experience a fitness cost when dietarily preformed sources of PUFAs are absent, and exhibit dramatic reduction in growth and reproduction (19, 20), suggesting that outsourcing PUFA production to either primary or more basal producers is preferred. Earthworms, as part of Annelida, may contain the enzymatic repertoire in their genome to produce long-chain PUFAs (18), though this has not been specifically explored in Lumbricidae. Terrestrial and freshwater vertebrates are physiologically constrained by the limited availability of long-chain PUFAs, and cope by either adjust dietary foraging behaviors, or through adaptive radiations to increase synthesis capacity encoded in the genome (19, 21). For example, mammals lack Δ12– and Δ15-desaturases, preventing the potential to synthesize linoleic and alpha-linoleic acid, which are precursors to longer-chain FAs (22). Therefore, EPA, ARA and DHA are considered essential in the mammalian diet. However, PUFAs accumulate at higher trophic levels, starting from microalgae and bacteria as primary producers (23, 24). Marine bacteria and microalgae use PUFAs in membrane lipids to maintain sufficient membrane fluidity in high-pressure and low-temperature environments (2, 9).

### Linking microbial activity to the food web

As PUFAs are important to human and animal development, physiology, and health, it is crucial to better understand PUFA producers and their role in the food web. This knowledge is particularly valuable to the biochemical engineering industry, as it aims to develop biosynthetic methods for PUFA production (25). Additionally, there is a significant goal to increase PUFA yield from natural microorganisms, reducing dependence on marine resources and promoting sustainable production (26). However, the supply of PUFAs is linked with aquatic organisms, and there is little evidence or expectation for a de-novo PUFA producer in terrestrial habitats, leaving open the question as to how these habitats are adequately supplied. Therefore, understanding how PUFAs are produced in terrestrial ecosystems and supplied to terrestrial food webs has cross-disciplinary interest from fields such as human evolution, ecology, and microbiology. The bacterial contribution to this effect has received relatively far less attention than have plant and eukaryotic producer systems, yet there is promising genetic and functional evidence that soil bacteria are capable PUFA producers (27). One unique study focused on the earthworm *Lumbricus terrestris*, and revealed increasing concentrations of PUFAs, including ARA, EPA, and DHA, from the surrounding soil to the intestinal contents – referred to hereafter as “gut-soil” – of *L. terrestris* (28). The authors concluded that a variety of unique fatty acids, including the long-chain PUFAs, are produced by microorganism inside of the earthworm gut, and that these lipids are then transferred from the intestine to the luminal cells and then finally into muscle tissue. The long-chain PUFAs were hypothesized to be used for energy needs of the *L. terrestris*. Another study analyzed the fatty acid composition of gut-soil from the epigeic composting earthworm, *Eisenia fetida*, in a comparison between bulk soil containing the earthworms, and soil without (29). The study reported a lower total fatty acid content in soil with *E. fetida*, along with reduced microbial biomass and diversity. The identified fatty acids ranged from twelve to twenty carbon atoms in length, but no unsaturation was detected among the twenty-carbon chain varieties. Overall, these studies report that earthworms modify soil properties and the microbial communities by selectively filtering microbial biomass, and additionally that earthworms likely benefit from the absorption of bacterial metabolites, including certain lipids.

Efforts have been dedicated not only to the exploration of fatty acid production in earthworms but also to the analysis of microbial compositions within diverse earthworm species towards understanding their role in composting and soil revitalization. One study examined the gut microbiome of earthworm species from three genera: *Aporrectodea, Allolobophora*, and *Lumbricus*, which occupy two different ecotypes. The *Lumbricus* are vertically burrowing “anecic” worms whereas the *Aporrectodea* and *Allolobophora* are horizontally burrowing “endogeic” worms (30). By utilizing 16S rRNA amplicon sequencing, it was discovered that Pseudomonadota was the primary phylum present in both earthworm ecotypes. Furthermore, the genus *Arthrobacter* and the class of Gammaproteobacteria were indicator taxa for *Lumbricus*, while at higher taxonomic levels the Acidobacter and Alphaproteobacteria were indicative of *Aporrectodea* (30). Further distinctions between ecotypes included presence of bacteriovore protists, as well as richness of bacterial taxa, which is considered to reflect the variation in feeding behavior of these earthworms. Another study focused on the top-soil and leaf-litter dwelling “epigeic” *E. fetida* and assessed the microbial composition of its microbiome when fed with different compost types, including brewers spent grain, cow manure, and a 50/50 mixture of both (31). Analysis of the earthworm castings (feces) revealed that, regardless of the treatment, the dominant phyla remained Proteobacteria and Bacteroidetes. However, a significant increase in Firmicutes and Actinobacteria was observed when *E. fetida* was fed with either 100% or 50% brewers spent grain. Similar results have been reported when *E. fetida* was fed vegetable waste (32). In sum, the effects of feeding behavior and the impact of soil conditions have been well studied in how they can modify composition of the earthworm gut-soil microbiota. These works lend important insights about how the earthworm gut-soil profile is distinct but related to the microbial composition of bulk soil. Variations between different types of earthworms appear dependent upon their ecotype rather than species.

These previous works suggest that the gut-soil microbiome of earthworms may contain PUFAs that are not initially present in the soil that they inhabit, and so, are not taken-up from primary consumption. However, the combined study of the gut microbiota and the lipid metabolites from the same sample set is needed to resolve the source of PUFA production and to define the microbial organisms involved in this production. We address this gap with an analysis of the fatty acid and microbial compositions of gut-soil, bulk soil, and earthworm castings of *E. fetida*. We hypothesize that bacteria in terrestrial environments, including those in the guts of terrestrial animals, may be capable of producing PUFAs, but only under certain conditions or with specific microbial neighbors. To test this hypothesis, the gut soil content of *E. fetida* was analyzed in three parts. First, the fatty acid composition was analyzed using gas-chromatography mass spectrometry (GC-MS). Second, the prokaryotic and eukaryotic constituents were profiled using 16S and 18S rRNA gene amplicon sequencing. Finally, amplicon sequencing was performed on the initiator ketoacyl-synthase (KS) domain on the polyunsaturated fatty acid gene (*pfaA*) (33) in order to identify microorganisms with the genetic potential to de-novo produce PUFAs using the iterative type-1 polyketide-like synthase system. This work uses the earthworm and its gut microbiota as a model for terrestrial ecosystem dynamics to identify potential PUFA producers and to help us understand the origin of PUFAs in terrestrial ecosystems and the diversity of microorganisms that may provide them.

## Results and Discussion

We analyzed the total fatty acid content and microbiome profiles of three sample types: pre-compost (PC), vermicompost (VC), and the gut soil (GS) of the earthworm *E. fetida.* PC is a mixture of horse manure and thermophilic compost used as feed and applied once per week, while VC is worm castings (feces) that is normally collected via sifting and used as an organic fertilizer. The sampling was conducted over two weeks, with samples collected on three specifically spaced days within each week. PC was only sampled on the first day of each weekly “feeding” cycle, as it is laid freshly onto the existing *E. fetida* compost housing once a week.

### Earthworm gut-soil harbors a distinct fatty acid profile enriched in PUFAs

The fatty acid profile of samples collected in week one and week two were compared to identify any batch effects between sampling weeks based on sample type or sampling days (as shown in Materials and Methods). Permutational multivariate analysis of variance (PerMANOVA) showed a significant difference (p = 0.016) between the sampling weeks, but this only explained 0.5% of the variation, while most of the variation (87.6%, p = 0.001) was explained by the sample type (Figure 1). We then examined whether there were any significant differences between the respective days of sampling for each sample type using the Kolmogorov-Smirnov test. No significant differences were identified for all sample types, therefore, the data from week one and week two were merged for all subsequent fatty acid composition analysis.

**Figure 1.**
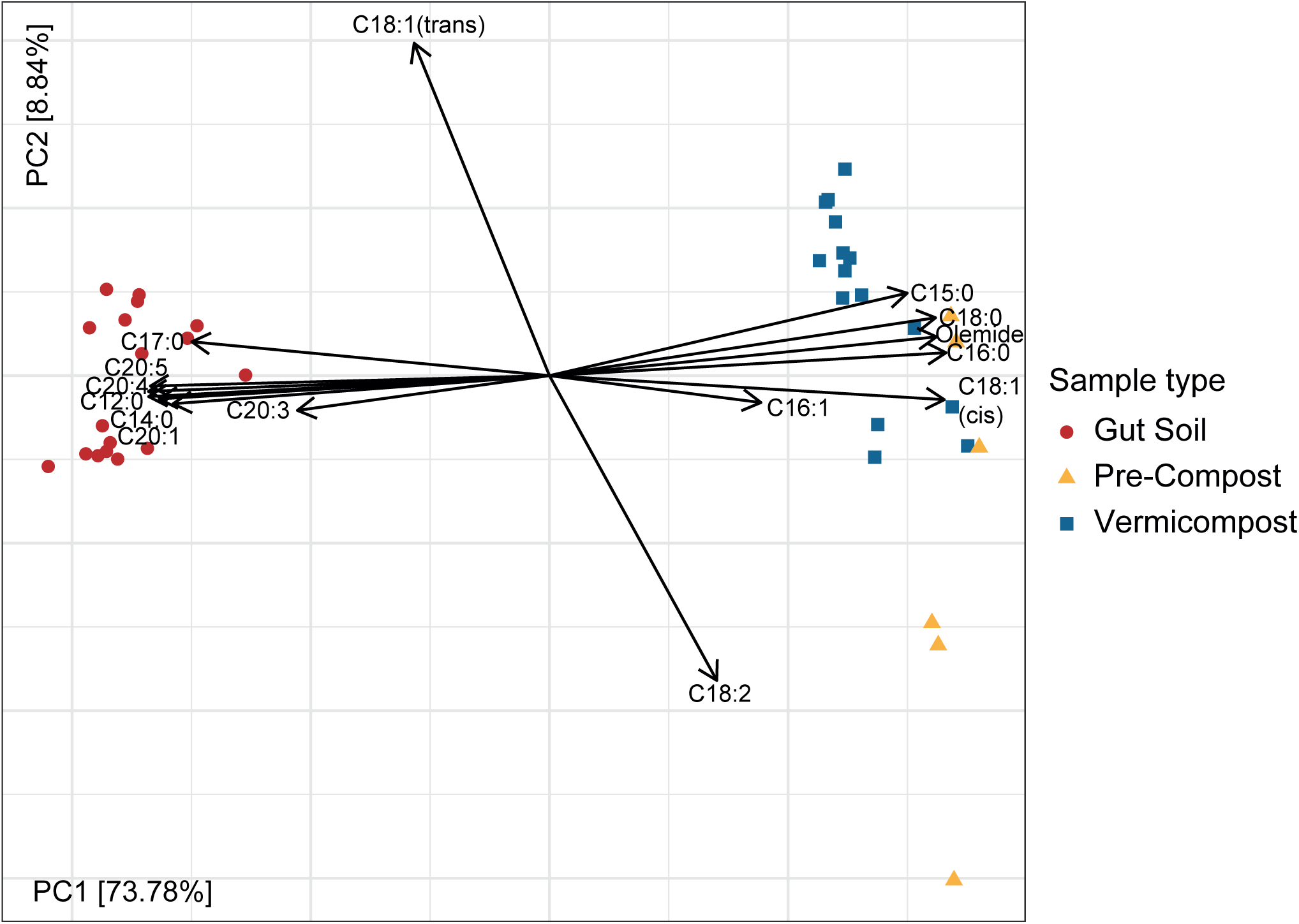
Principal component analysis (PCA) of the fatty acid content extracted from pre-compost (yellow triangle), gut soil (red dot) and vermicompost (blue square) after center-log-ratio transformation (CLR) with each loading arrow representing one fatty acid identified.

According to the analysis of the fatty acid composition of PC presented in Table 1, palmitic acid (16:0) was found to be the most abundant fatty acid, constituting on average over 30% of the detected fatty acids, followed by stearic acid (18:0) and oleic acid (18:1 *cis*-ω9) at 13% and 14%, respectively. The only PUFA detected in PC was linoleic acid (18:2 ω6), which accounted for more than 6% of the total fatty acid content, and likely derives from bulk plant material in the PC mixture. Linoleic acid is known to be a precursor for the synthesis of the omega-6 series of fatty acids, particularly leading to ARA, whereas α-linolenic acid, which is the precursor to the omega-3 series, including EPA, and DHA, was not detected.

**Table 1.**
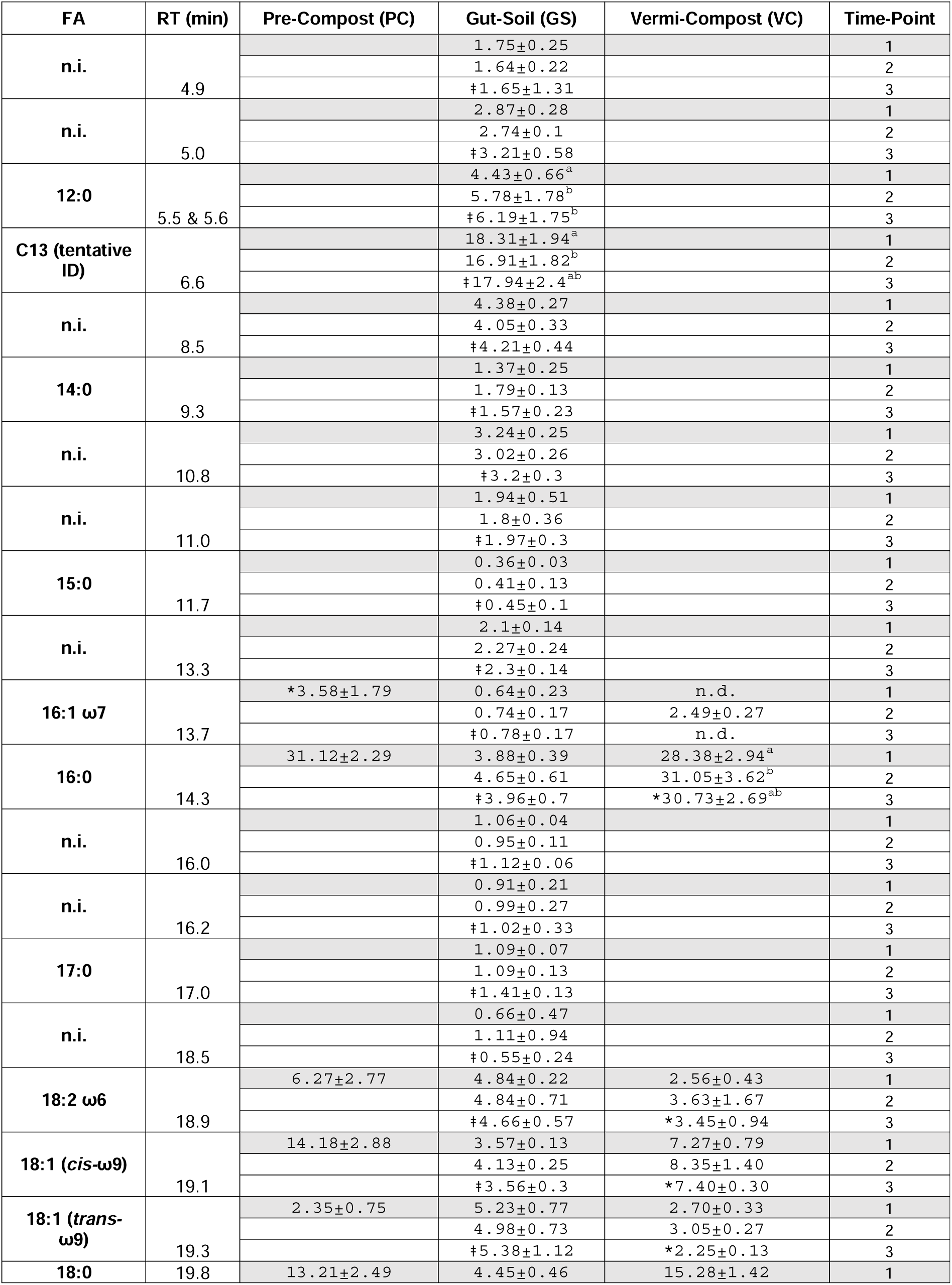
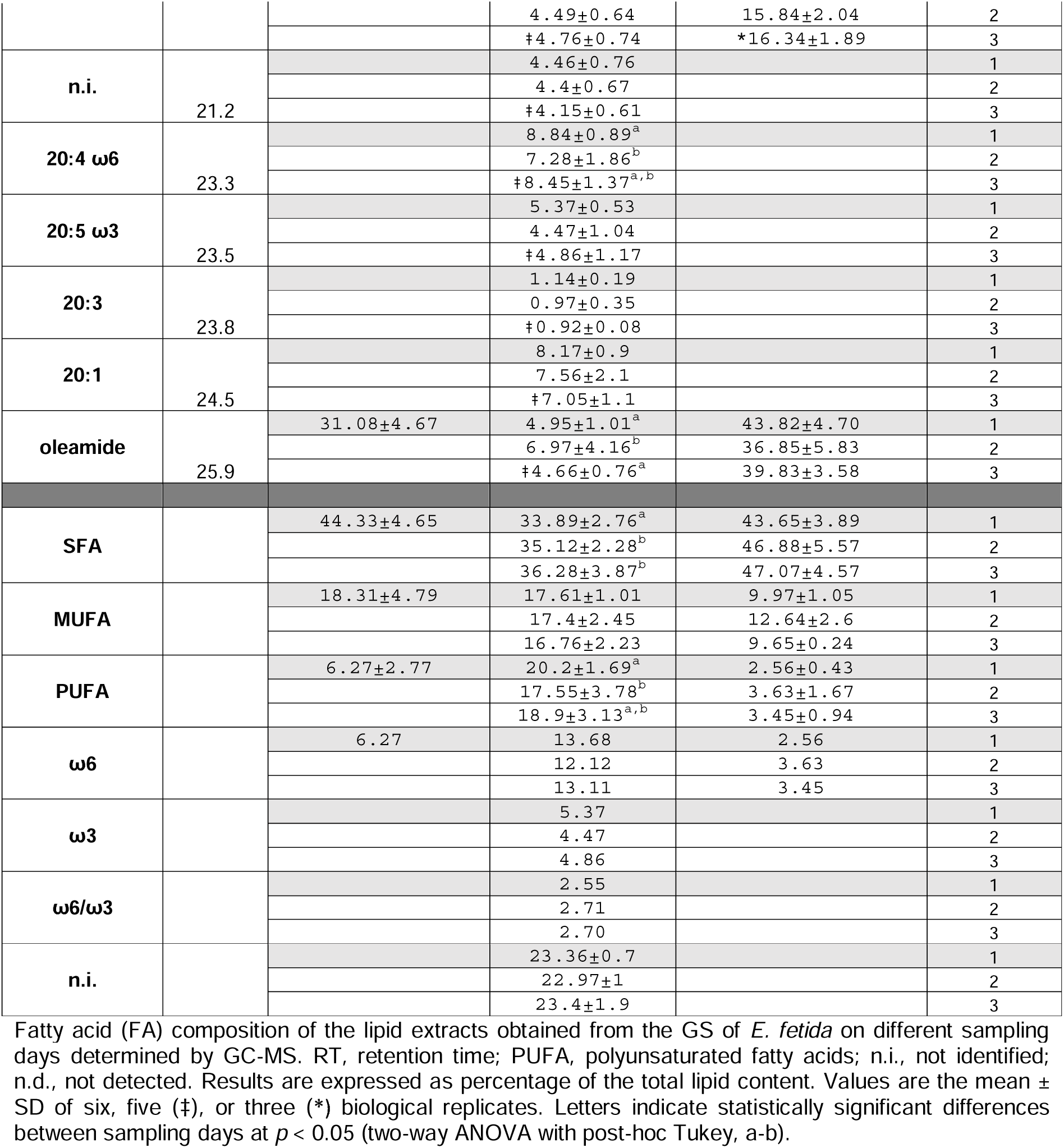
% Fatty Acid of Total Lipid Content ± SD.

One curious finding was the presence of *trans*-fatty acids. Although bacteria do not usually produce *trans*-fatty acids, they can produce them under specific circumstances, which has been demonstrated in *Vibrio* and *Pseudomonas*. In *Vibrio*, a temperature shift caused cis/trans isomerization, while in *Pseudomonas*, toxic compounds such as organic hydrocarbons resulted in isomerization (34–37). PC partly consists of thermophilic compost which has been heat processed through fermentative activity, and this temperature increase could potentially have caused *cis/trans* isomerization from bacterially originated FAs.

The percentage of PUFAs in GS was found to range from 17 to 20% of all FAs across the three sampling days. ARA (20:4 ω6) and EPA (20:5 ω3) were identified as two notable fatty acids, comprising ∼7-9% and 4.5-5.5% respectively of the detected fatty acids. Other major fatty acids included lauric acid (12:0, ∼4-6%), elaidic acid (the trans isomer of oleic acid,18:1 *trans-*ω9, ∼4-6%) and gondoic acid (20:1, ∼6-9%). No significant differences in the relative abundance of fatty acids were observed between sampling days (p > 0.05), except for lauric acid (12:0) (two-way ANOVA with post-hoc Tukey, day 1 – day 3: *p* = 0.0280), arachidonic acid (20:4 ω6) (two-way ANOVA with post-hoc Tukey, day 1 – day 2: *p* = 0.0451), the overall SFA content (two-way ANOVA with post-hoc Tukey, day 1 – day 2: *p =* 0.0002, day 2 – day 3: *p* = 0.0002) and the overall PUFA content (two-way ANOVA with post-hoc Tukey, day 1 – day 2: *p* = 0.0002). PUFAs were found to have a lower relative content on day one of sampling compared to day two. Lauric acid content increased steadily from day one to day three, but only the difference between day one and three was found to be significant. This may be due to the feeding cycle, which occurs on day one of the week, and therefore the second sampling point captures the peak nutrient availability and metabolic activity, which then falls off by the end of the week. The high percentage of unidentified fatty acids (n.i.), which accounted for approximately 23% of the total content, were compounds either structurally similar to fatty acids but unidentifiable with the standard fatty acid methyl-ester (FAME) mix, or compounds with low identification scores in the national institute of standards and technology (NIST) mass spectral library and thus grouped together as “not identified”.

Only six fatty acids were identified in VC, all of which were the same as those found in PC samples. Among these, linoleic acid (18:2 ω6) was the only PUFA, accounting for less than 4% of fatty acids. Palmitic acid (16:0) was one of the major fatty acids found in VC, accounting for 28-31% of the total content, along with stearic acid (18:0), which accounted for approximately 15%. Palmitic acid showed significant differences over the one-week sampling period (two-way ANOVA with post-hoc Tukey, p = 0.0321). We detected oleamide, or oleic acid amide, in all samples, which likely originated from the compost added to PC, since oleamide has been detected in mixtures containing starch, sunflower oil, and soy protein (38). These results show that PUFAs are uniquely concentrated in GS samples compared to the starting compost and the resulting vermicompost (Figure 1).

These results are largely consistent with the findings of Sampedro et al. (2006), who reported elevated levels of PUFAs including ARA, EPA, and DHA in the gut of *L. terrestris* compared to the surrounding soil (28). In our study with *E. fetida*, the PC is the nutrient rich housing soil for the metabolic activity of gut-resident microorganisms, the products of which are available for host uptake. Since we did not find PUFAs in PC (incoming) nor VC (outgoing) samples, then PUFAs found in the GS, either from microbial activity or from the shedding of earthworm epithelial cells, appears to be selectively retained in the earthworm and its microbiome for physiological needs. The diverse range of fatty acids that we recovered in the GS may be indicative of high metabolic activity stimulated in the gut.

### Metataxonomic amplicon analysis

To identify potential microbial producers of ARA and EPA in the gut soil samples, we used 16S and 18S rRNA gene sequencing alongside amplicon sequencing of the PfaA-KS domain using primers previously developed to target the PUFA synthase complex (described in the next section) (33). By using high-level taxonomic binning of the functional gene sequences and then examining members of these taxonomic bins among the sample types, potential producers of ARA and EPA could be identified. To account for effects of low biomass and interindividual variation, samples of GS from individual *E. fetida* (denoted IG) were also compared with the pooled GS samples (see Table S1 for sample metadata).

The V4 hypervariable region of the 16S rRNA was targeted using the 515F/806R universal primers with slight modification, and the TAReuk454FWD1/TAReukREV3mod universal primers were used to target the eukaryotic 18S rRNA gene for amplification and sequencing on an Illumina MiSeq (see Materials and Methods for primer sequences). After decontamination of 16S rRNA sequences, an average of 8116 (±4193 SD) reads were obtained for GS, 5606 (±4124 SD) for IG, 8296 (±5664 SD) for PC, and 7648 (±4295 SD) for VC. A total of 4238 ASVs were identified across all the samples. The 18S sequencing resulted in an average of 30753 (±15703 SD) reads for GS, 25977 (±10907 SD) for IG, 18221 (±12272 SD) for PC and 20034 (±14504 SD) for VC after decontamination with a total of 886 identified ASVs (see Tables S2 & S3 for 16S and 18S ASV tables).

### Prokaryote taxonomic sequence profiles

Observed ASVs and Shannon metrics were used to analyse alpha-diversity of the microbiome, based on a rarefied ASV table with 5000 reads for 16S rRNA analysis and 10000 reads for 18S rRNA analysis (Figure S1 rarefaction curves). No significant difference was found in richness or diversity between GS and IG samples. However, gut soils (GS & IG) showed significant variation compared to compost samples (PC & VC), with VC displaying significantly higher measures of richness compared to both gut soils (Wilcoxon, GS-VC: observed ASVs p = 3.291e-04 and IG-VC: p = 1.842e-05; Figure 2a). Similar variation was observed in Shannon diversity metrics, with significant differences identified between both gut soil samples compared to VC (Wilcoxon, GS-VC: Shannon p = 0.001636 and IG-VC: p = 9.578e-06; Figure 2a). This suggests that VC and PC have a higher evenness and contain more ASVs that were only identified once or twice compared to GS samples. Beta-diversity analysis was conducted using the unrarefied ASV table, but was normalized using a variance stabilizing transformation with DESeq2. Ordination analysis using Euclidean distance on center-log-ratio transformed data indicated distinct sample clusters for PC and VC, while IG and GS clustered together (Figure 2b).

**Figure 2.**
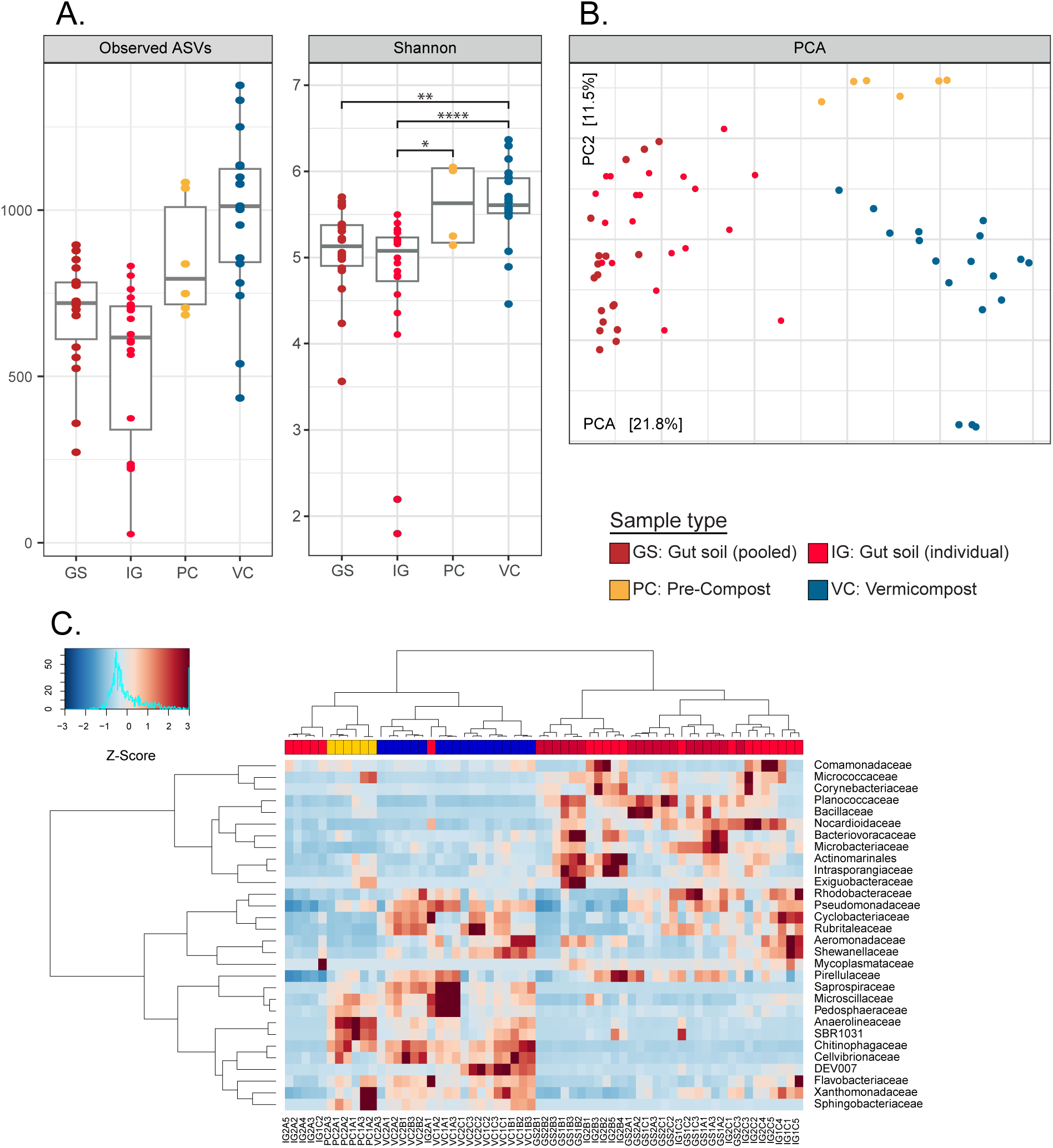
Alpha– and beta-diversity analysis comparisons for gut soil (GS, dark red), individual gut soil (IG, bright red), pre-compost (PC, yellow) and vermicompost (VC, blue). (**A**) Boxplots for Observed ASVs and Shannon alpha-diversity metrics with significant differences shown for different sample types with pairwise comparisons between groups (p > 0.05 (ns), p ≤ 0.05 (*), p ≤ 0.01 (**), p ≤ 0.001 (***), p ≤ 0.0001 (****)). (**B**) Ordination of ASVs using Euclidean distance metrics shows separation of sample types. (**C**) Ward’s clustering of rarefied ASVs binned at family-or lowest assigned taxonomic level and filtered for 2% abundance in at least 5% of samples.

Based on the analysis of alpha– and beta-diversity, it was found that there were no significant differences in richness or diversity between the pooled and individual GS samples, and that they clustered together. In contrast, the PC, VC, and gut soil samples were found to each have distinct microbial community compositions. Figure 2c shows clustering of class-level bins and the relatively enriched or depleted taxa across samples, and reveals that GS samples largely cluster together except for a few IG samples, while the compost samples form a separate cluster. This suggests that either the earthworm gut community is truly an endemic microbiome, or that the gut environment provides highly selective conditions that promote transient growth and activity in a stable reproduceable manner (39).

The relative abundance of major phyla and families in each sample type provides additional evidence of different microbial community compositions per sample type. Phyla that contribute at least 5% and families that contribute at least 2% to the total relative abundance are compared for relative abundance. To test for significant differences in relative abundance, a pairwise Wilcoxon analysis with post Benjamini-Hochberg was conducted between each sample type (Figure 3, and see Figure S2 *pairwise family level corr*). Comparing IG and GS samples revealed significant differences across sample types at the phylum, class, and family level, however, these differences were primarily driven by only a few family-level groups within the primary segregating phyla. Specifically, significant differences were found in three of 51 families in Gammaproteobacteria, five of 61 families within Firmicutes, and seven of 37 families within Bacteroidota for gut sample comparisons. On the other hand, pronounced differences were observed in the comparisons between the gut soils and composts, where most families showed significant changes in abundance. In general, there was significant variation in relative abundances when gut soils were compared with composts (Figure S3a *relative abundance barplots*). While the significance was less for IG than for GS in most cases, the trend of the mean values was the same, supporting that the features seen in pooled samples can be generalized to individual specimens, and are not driven by stochastic or random features from outliers.

**Figure 3.**
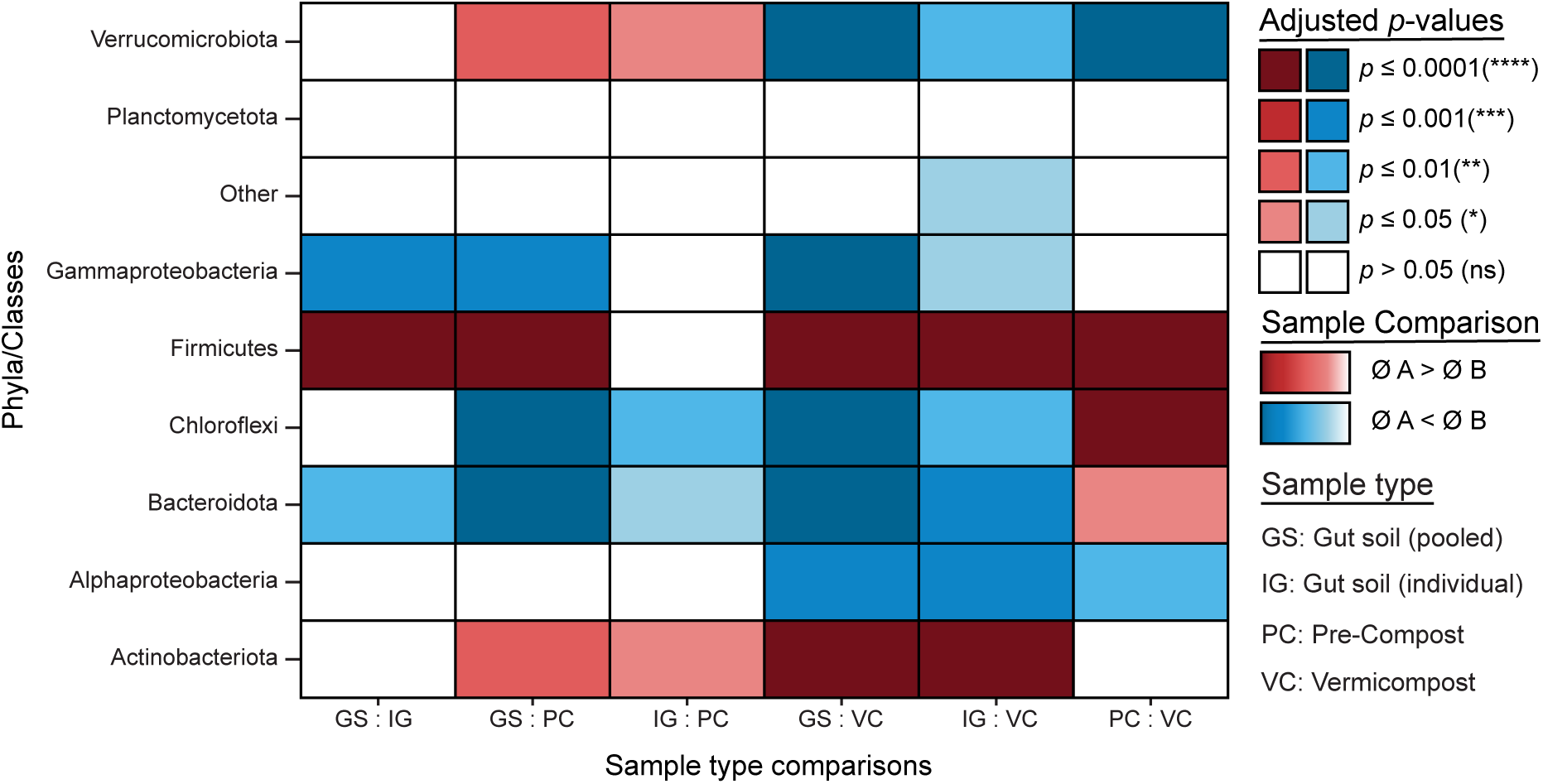
Pair-wise comparisons of each sample type by Wilcoxon test with Benjamini-Hochberg corrected p-values for each comparison. (p > 0.05 (ns), p ≤ 0.05 (*), p ≤ 0.01 (**), p ≤ 0.001 (***), p ≤ 0.0001 (****)). Blue color scale indicates that the mean of sample A is lower than in sample B. Red color scale indicates that the mean of Sample A is greater than the mean of sample B. Gut soil (GS), individual gut soil (IG), pre-compost (PC), vermicompost (VC).

Notably, Actinobacteriota and Firmicutes display a higher relative abundance in gut soil samples compared to the compost samples, whereas Bacteroidota show a higher relative abundance in the compost samples compared to the gut soils.

The pair wise comparison of sample types supported the findings from the taxonomic summary regarding Actinobacteriota, Bacteroidota, and Bacillota (formerly Firmicutes). Specifically, Bacteroidota, Gammaproteobacteria, and Chloroflexi had a higher relative abundance in both compost samples compared to each of the gut soil samples. This was also true for Alphaproteobacteria when comparing the gut soil samples to VC. Verrucomicrobiota had a higher relative abundance in the gut soils compared to PC, but still lower than in VC. No differences were found in Planctomycetota for all comparisons and in Alphaproteobacteria when comparing the gut soils to PC.

In previous works, the major phyla identified in *E. fetida* when fed with different types of compost were Proteobacteria and Bacteroidota. However, since the compost used in our study differed from that used in the analysis by Budroni et al. (2020), slight differences in the microbial composition of GS are expected (31). This is supported by previous reports that show changes in microbial composition based on the compost composition, as well as other studies that varied the feeding compost (40). A study by Sapkota et al. (2020) also identified Proteobacteria as a major phylum in their analysis for the earthworm genera *Aporrectodea* and *Lumbricus* (30). Although Proteobacteria and Bacteroidota are still present in GS from our study, they do not account for most of the microbial composition in this sample type. Their lower abundance could be explained by the nutrient-rich PC substrate, applied to the earthworm population as feed and housing matrix. Previous research has shown that Proteobacteria and Bacteroidota are particularly increased in abundance when *E. fetida* is fed with a nutrient-poor substrate, whereas a nutrient-rich substrate results in higher microbial diversity and a higher abundance of Firmicutes and Actinobacteria (40).

### PfaA-KS for assessment of bacterial PUFA producers

To understand the potential that PUFA lipids concentrating in the gut of the earthworm are derived from bacteria, we used a previously developed universal primer targeting the beta-ketoacyl synthase (KS) region in the single-copy pfaA gene, which amplifies an approximately 502 bp region spanning the PfaA-KS N– and C-terminals (33). The KS domains from PKS types tend to cluster as a monophyletic group, and within these groups, the KS can approximate the evolutionary phylogeny of the organisms (41, 42). In addition, the KS sequence phylogeny is distinctive of the type of PKS complex in which they occur, which makes it a suitable candidate marker for targeting the genes involved in synthesis of a specific metabolite. We sequenced the amplified KS products using an Illumina MiSeq, resulting in 53 samples with successfully generated, filtered, and PfaA-KS-assigned reads, and final average sequence counts segregating per sample type: 16 (±27 SD) for GS derived reads, 315 (±526 SD) for PC, and 1230 (±1459 SD) for VC. A total of 66 PfaA-KS ASVs were derived using the DADA2 workflow on merged reads. One read (Seq64) was probably a chimera from earthworm DNA and therefore removed from the dataset before proceeding to downstream analyses. The final 65 PfaA-KS ASVs led to the identification of 6 resolved unique bacterial strains within 14 taxonomic bins across 5 phyla (Table S5). Neither microeukaryotes nor fungi were found among the annotations, although some of these organisms harbor an iterative type I PKS (T1PKS) gene complex and KS in a neighboring clade to bacterial iterative T1PKS (15, 42, 43).

Little is known about the distribution of PUFA-synthase genes and production potential in microorganisms outside of marine contexts, with only very recent investigations beginning to probe the terrestrial biospheres (17, 44). To date, we are aware of only one primer set that has been developed for specifically targeting the ***pfa*** T1PKS KS domain, which was modeled from the sequences of marine Pseudomonadota, and especially those belonging to γ-Proteobacteria. Since these bacteria appear to have propagated T1PKS genes via HGT (42), a single primer pair may work universally within this phylum to capture many ***pfa*** KS sequences. However, this specificity towards marine Gammproteobacteria likely hinders recovery of bacteria in diverse phyla deriving from terrestrial soil and freshwater ecosystems such as actinobacteria and cyanobacteria respectively, where T1PKS appear to have evolved rather from a common ancestor. Indeed, certain soil taxa with known PUFA production competence were surprisingly not found among our annotated sequences, such as some strains within myxobacteria, Actinomycetes, and Desulfobacteria (27, 45–47). We checked primer specificity to 427 phylogenetically diverse organisms from NCBI GenBank reference genomes and saw alignments occur across eight deeply divergent lineages, including common soil or environmental bacteria. Gammaproteobacteria (n=151) were the dominant lineage, of which *Shewanella* (n=107) genomes were most highly represented. Of 427 queried genomes, 182 matched both forward and reverse primers and indicated high likelihood of successful amplification. This analysis did confirm that *Sorangium cellulosum* and *Minicystis rosea* of Myxococcota would be missed (Figure S4 *Phylogenetic distribution of PfaA-KS primer matches*). Despite this limitation, a surprising number of sequences and diversity were recovered from the compost and earthworm gut soil sample material that we report here. We recovered PfaA-KS sequences that had >95% sequence identity to those found among marine bacterial reference isolates using BLASTn query on the full nucleotide collection (nt). For sequences that were not well resolved using the NCBI RefSeq nt database, a BLASTx translated amino acid sequence was mapped against the non-redundant (nr) database, with most hits achieving >95% homology (see Tables S6 and S7 for a full summary of BLAST hit results).

### Phylogenetic inference among recovered sequences

In order to understand the relationship of the sequences we captured from earthworm gut and compost soil samples, we constructed a phylogenetic inference tree using our 65 bacterial PfaA-KS ASVs plus the 446 PfaA-KS sequences that were published in the original publication for creation of the PfaA-KS primer set (33) (Figure 4; Table S8). The relevant difference with our present study is use of the PfaA-KS primers on terrestrial soil-derived samples, whereas prior research has been restricted to their application on marine water and sediment samples. Using the *Escherichia coli fab*F gene as an outgroup (representative of a KS subtype involved in the canonical type II FAS system), we see that our sequences surprisingly form clusters among shallow branches of the tree which match their taxonomic assignments, rather than their environmental source. However, known evolutionary relationships that are based on whole-genomes are not recapitulated, since more distantly related taxa (such as Pseudomonadota and Bacteroidota) form larger clusters that break up established evolutionary lineages, such as between members of Gammproteobacteria.

**Figure 4.**
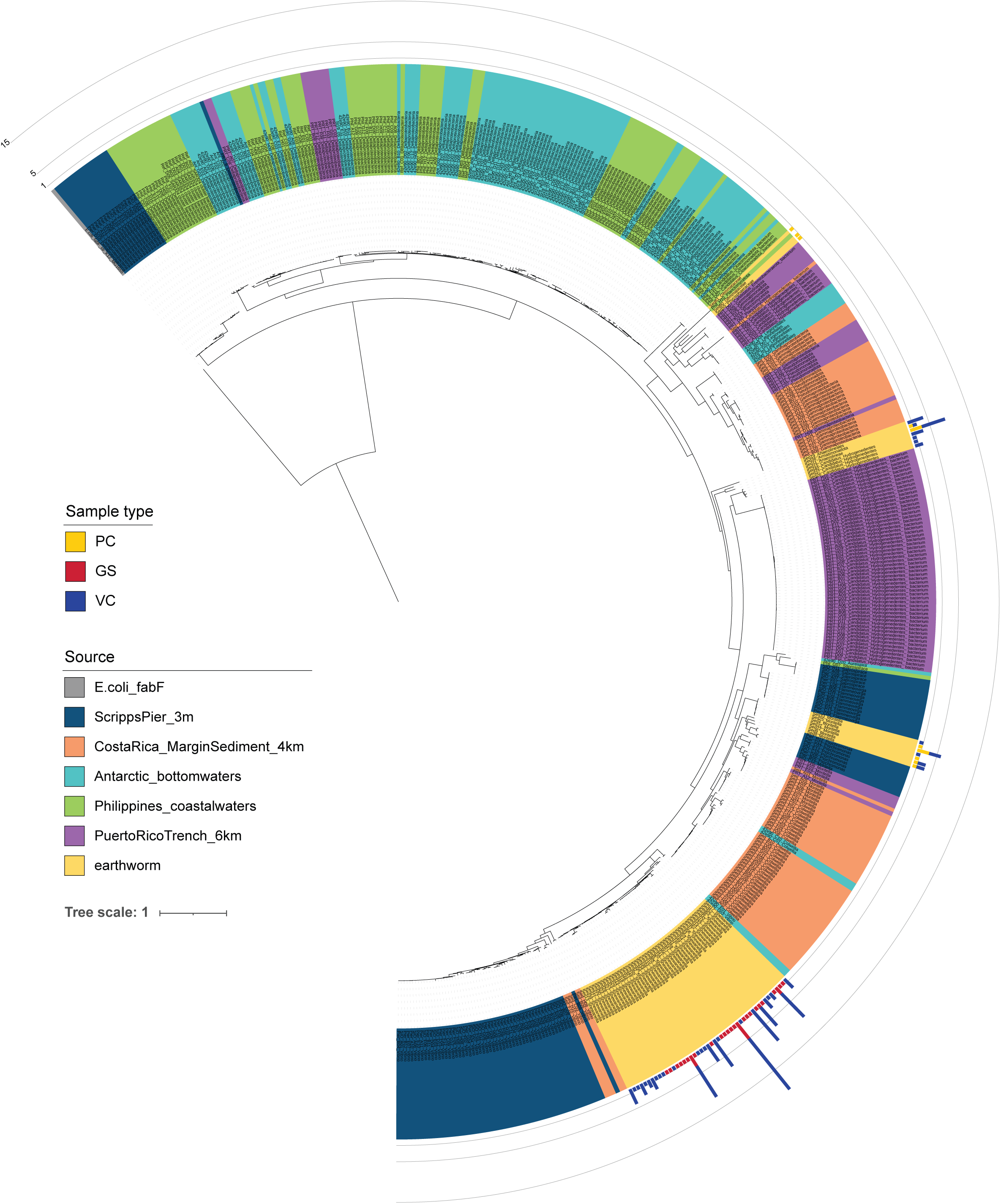
Phylogenetic inference of 511 PfaA-KS amplicon sequences from this study (earthworm) and prior work investigating marine bacterial sources of PUFA synthesis. The *fabF* gene from *E. coli* was used as outgroup and root. Stacked barplots show the number of samples that contain the ASV sequence and are colored by the sample type.

There is a clear separation between sequences found within earthworm GS samples and those of the compost samples (denoted by barplots along the outer ring of the tree). All GS sequences are assigned to *Shewanella* and cluster among a fairly undifferentiated lineage that include other *Vibrio* assigned sequences, and which stem from a bifurcation that separates 1) facultatively anaerobic cold-water-marine Gammaproteobacteria (*Vibrio*, *Colwellia*, *Moritella*, *Psychromonas*) and aerobic saprophytic or predatory Bacteroidota (*Flammeovirga*, Saprospirace); from 2) subsurface anoxic dwelling taxa (Legionellales, Candidatus Hydrogenedentes), anaerobic fermenters (*Anaerolineae*) and terrestrial aerobic chemoheterotrophs (Armatimonadota) (48–50). The PfaA-KS sequences found within compost samples were spread more widely across the tree among organisms such as *Gemmata* (Planctomycetota), Cloacimonadota, and other Anaerolineales that are often found among carbon-rich sediments such as peatbogs, waste-water treatments, or the rhizosphere. To date, the activity and product of these putatively competent PUFA synthase complexes remains unconfirmed (47). The remaining one-third of the tree is wholly dominated by a basal cluster of sequences assigned to Deltaproteobacteria or its sublineage, SAR324, which are distinguished from the other taxonomically and phylogenetically diverse branchings. This suggests that metabolic niche participation has broadly determined the structure of the tree, which separates primarily between marine chemolithoheterotrophs (sulfate reducers) (51) and diverse chemoorganoheterotrophs (carbon/nitrogen utilizers through predatory or saprophytic processes) (48, 52). This topology further supports the observation that T1PKS are prone to HGT events (42), and that HGT propagation of the *pfa* complex helps to explain its environmental and phylogenetically widespread occurrence and lack of strict coherence among species or sub-species level presence/absence patterns (47).

### PfaA-KS sequence distribution and abundance across sample types

To gain insight about which members of the microbial community that are concentrated in earthworm gut-soil and which may be contributing to production of PUFA, we conducted in-depth investigation on the ASV table abundances, which was first transformed to center-log-ratio (CLR) values (53). An extremely sparse compositional table rendered normalization practices such as rarefaction and relative abundance inadequate to standardize sequence abundances across samples. We confirmed that there is no apparent bias in the amount of genetic data recovered from the samples based on sample type, starting amount, or concentration, supporting the conclusion that the observed sparsity of the PfaA-KS ASV table is biologically real (see Table S1 for sample metadata; see Figure S5 for associations between PfaA-KS sequence recovery and sample metadata). Surprisingly, in contrast to the metabolite lipid profiles generated from the same sample material, the recovered PfaA-KS sequences were most diverse and concentrate among compost samples (PC and VC) rather than the GS samples (Figure 5a). The PfaA-KS sequences were sparsely distributed at very low abundance in GS samples, both pooled and individual (Figure 5b). While the ASV taxonomic assignments suggest that just a handful of unique taxa contribute to the total sequence variation, the sequence variants themselves are quite numerous even within the same putative bacterial strain, and are strongly segregated between GS and compost samples (Figure 5c). No single ASV is shared across all three sample types. This can be most clearly seen when ASVs are aggregated at genus level, or otherwise their lowest taxonomic bin, and then hierarchically clustered according to Ward’s sum of squares method (Figure 5b). The GS samples are composed entirely of *Shewanella* assigned sequences while the compost samples contain a mixture of the other seven environmental taxa bins. This suggests that only *Shewanella* are remaining in close association to the earthworm host tissue rather than passing through as transients (indicated by the strains found in VC but not in GS), and perhaps actually are co-residents of the earthworm gut (54).

**Figure 5.**
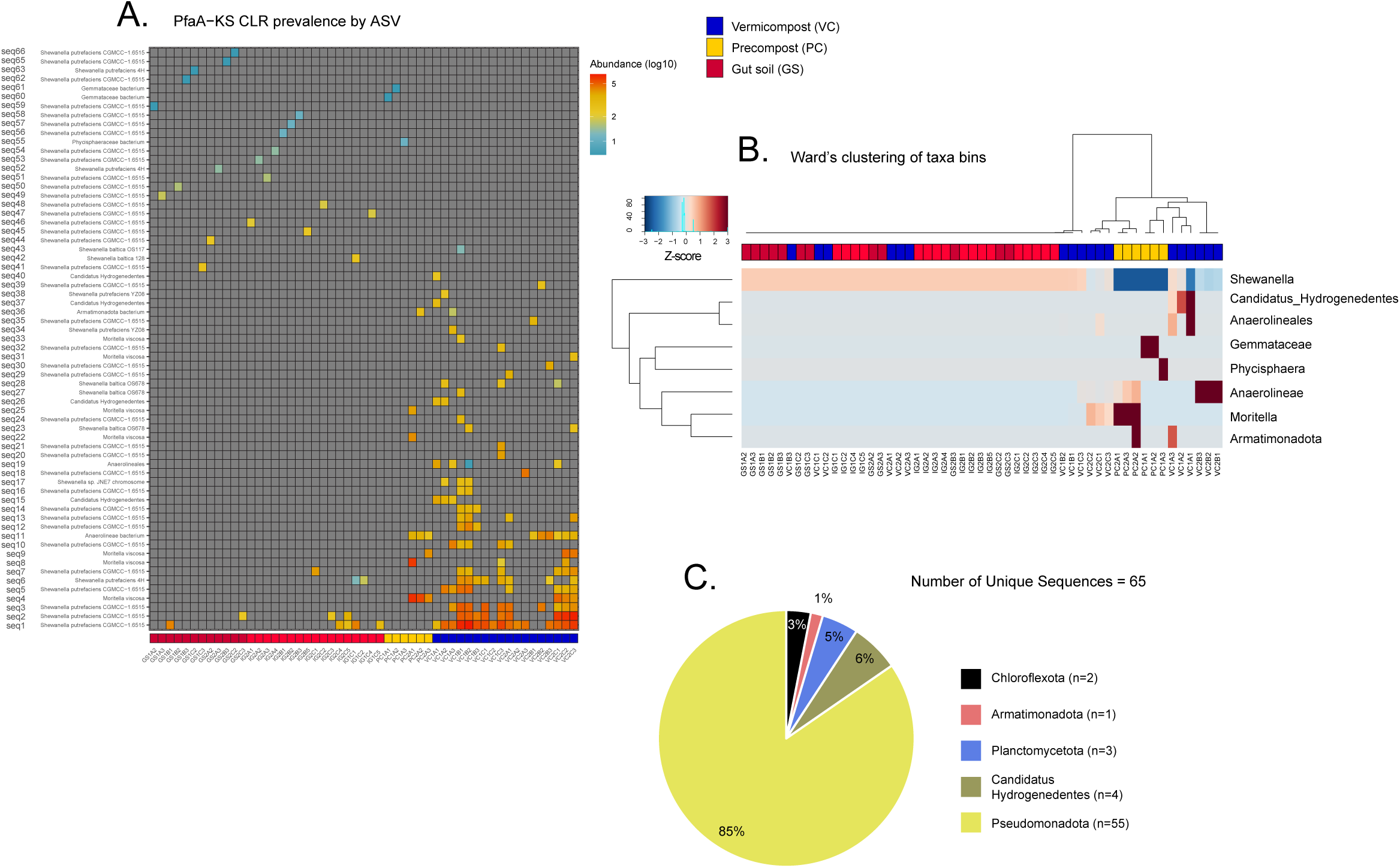
Visualization of the sparse distribution and low abundance of 65 bacterial PfaA-KS ASVs by sample type based on center-log-ratio transformed ASV table. (**A**); and hierarchical clustering of samples based on ASVs binned at lowest taxonomic assignment (**B**). Number of unique sequences shown across 5 phyla assignments shows clear dominance by Pseudomonadota, and primarily from the genus *Shewanella* (n=48) (**C**).

### PfaA-KS taxa within the microbiome profile

To understand the prevalence of putative PUFA-producing taxa among the overall prokaryote microbiome community, the taxonomic bins assigned to the PfaA-KS sequences were sought among the 16S rRNA dataset, and the matching ASVs from the abundance tables were subset for further analysis (Figure S6 *prok.pfa abundance barplots*). The subset of PfaA-KS taxa from the 16S data (hereon referred to as **prok.pfa**) were normalized by the total number of ASVs, and then transformed using CLR to compare with the PfaA-KS sequence amplicon dataset (hereon referred to as **pfa**). First, Procrustes analysis was used to infer the correspondence of the ASV level ordinations of the **prok.pfa** and **pfa** data using Euclidean distance. Using the function “protest” with 999 permutations, the datasets were found to have modestly significant correlation (m12 squared = 0.758, corr. = 0.492, p = 0.001), and their ordinations show similar separation of sample types along the first and second principal components (Figure 6a). Next, the ASVs counts were binned at genus level (or lowest taxonomic assignment), resulting in seven PfaA-KS taxonomic assignments: Anaerolineales, Armatimonadota, *Gemmataceae*, Hydrogenedentes, *Moritella*, Phycisphaera, and *Shewanella*. The correspondence of these binned taxa vectors within the **pfa** and **prok.pfa** abundance tables was calculated using a Mantel test on the Euclidean distance matrices, yielding again low correlation (Spearman’s rho = 0.194, p = 0.007, Figure S7 *mantel test*), suggesting that the abundance of the sequences captured using the PfaA-KS primers cannot be related to the microbiome profile reconstructed from 16S rRNA amplification. Correlation and visualization of the abundance distribution of each taxonomic bin shows that only for *Shewanella* abundance is there coherence between the PfaA-KS and 16S rRNA abundances (Pearson’s rho: PC = 0.465, GS = 0.662; Figure 6b & 6c; Figure S8 *TaxaLinearAssoc-pfaclrM2;* & Figure S9 *Histograms*), supporting a hypothesized situation that *Shewanella* proliferate in the earthworm gut relative to the overall community, during which they could contribute PUFA nutritionally to the earthworm, and are not just transient dead organic matter. To be able to truly correlate the amplicon datasets requires quantification of the amplified gene regions, and more targeted research in the future should incorporate a digital PCR or quantitative PCR step in the amplicon preparation workflow. This would verify the coherence in the presence of the PfaA-KS and the 16S gene regions and the bacterial taxa from which they derive.

**Figure 6.**
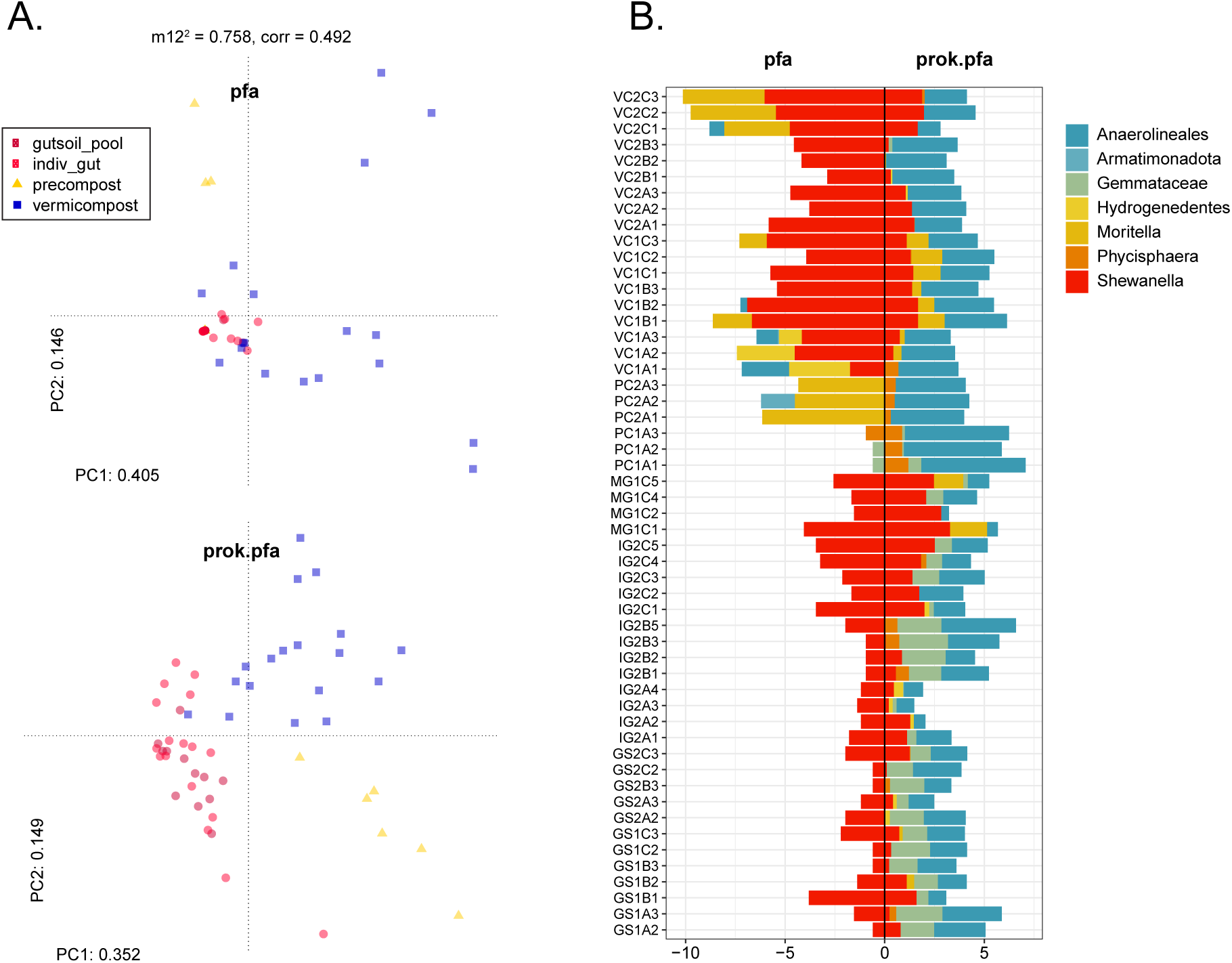
Comparison of ASV abundance tables for the PfaA-KS amplicons (**pfa**), and the subsetted PfaA-KS assigned taxa from the 16S rRNA data frame (**prok.pfa**). Ordination and Procrustes analysis on the transformed data frames (**A**) indicates modest correlation, shown by the sum of squares (m12^2^ = 0.758) and correlation coefficient (corr = 0.492). The principal component plots of the CLR-transformed data frames demonstrate this correlation from some interspersion of GS and VC samples, and with PC samples clustering separately. The abundance values of the binned taxa for **pfa** and **prok.pfa** show the distribution of taxa by sample (**B**).

Using effect-size as a measure of explanatory power for the difference in ASV abundance in the **prok.pfa** table also found Shewanella-assigned ASVs to be highly indicative of differences between vermicompost (earthworm castings), and pre-compost (compost soil applied as feed but not containing earthworms) (Tables S9 & S10; Figure S10 *Effect size aldex2 results*). The dissociation between **pfa** and **prok.pfa** abundances for the remaining taxa is most likely because the taxa that contain a putative PUFA gene complex are extremely low-abundant members of the total microbial community (especially in soil or compost), and so are negatively biased during 16S rRNA primer annealing and amplification.

### Eukaryote taxonomic assessment for other soil microbial or microfaunal PUFA producers

Earthworms ingest a variety of non-bacterial microbiota from the soil and can even be selective foragers. Phospholipid fatty acid (PLFA) biomarkers indicate the concentration of microeukaryotes and fungi within the earthworm gut relative to the bulk soil (55), and a robust immune response has evolved to entrap and expel parasitic nematodes (56, 57). Prior research on the lipid content of earthworm GS attributed the accumulation of PUFA in GS and somatic tissue to the metabolic activity of microeukaryotes such as fungi and protozoa. This was in accordance with the understanding that certain FAs are lipid biomarkers for the presence of organismal groups, on account of the FA composition of their phospholipid membrane (58). According to the pattern of observations, algae, yeast and other fungi are typically associated with synthesis of long-chain polyunsaturated acyl-moieties, including 20C+ fatty acids and sphingolipids, from both denovo and precursor 18C fatty acids (18, 28, 58–61). Common terrestrial bacteria on the other hand are attributed with medium-chain, straight or branched, odd-numbered-saturated or even-numbered-monounsaturated fatty acids (58, 62), and so they have not been considered candidate producers for PUFAs found in a terrestrial sediment context. Examples of soil-living microeukaryote organisms with notable concentrations of PUFA include invertebrate microfauna such as nematodes (Chromadorea), and macroinvertebrate collembola (Ellipura), fungi such as *Mortierellaceae*, and protozoan microalgae such as Labyrinthulomycetes. However, the origin of earthworm gut PUFAs from a microeukaryotic source has not been fully investigated. Therefore, we used 18S rRNA amplification to catalogue the presence of putative PUFA producing eukaryotes, and to determine whether their pattern of distribution across sample types may be a factor towards explaining PUFA accumulation in the earthworm gut environment. Eukaryote ASVs that mapped to the Oligochaetes subclass (aquatic and terrestrial worms) were first removed to eliminate the host signal that otherwise obliterated our ability to see representative taxa. For GS samples, this unfortunately resulted in a loss of >90% of reads. Filtered samples were then rarefied to 1000 reads and ASVs were binned at the “class/order/family” level for visualization and assessment (Figure 7, Figure S3 *Eukaryote taxa assignments barplot*). Of the resulting 46 taxonomic bins achieving at least 1% abundance in 5% of samples, four are identified as containing known PUFA producers: Chromadorea (nematodes), Ellipura (collembola), *Mortierellaceae* (soil fungi), and *Thraustochytriaceae* (saprotrophic protists).

**Figure 7.**
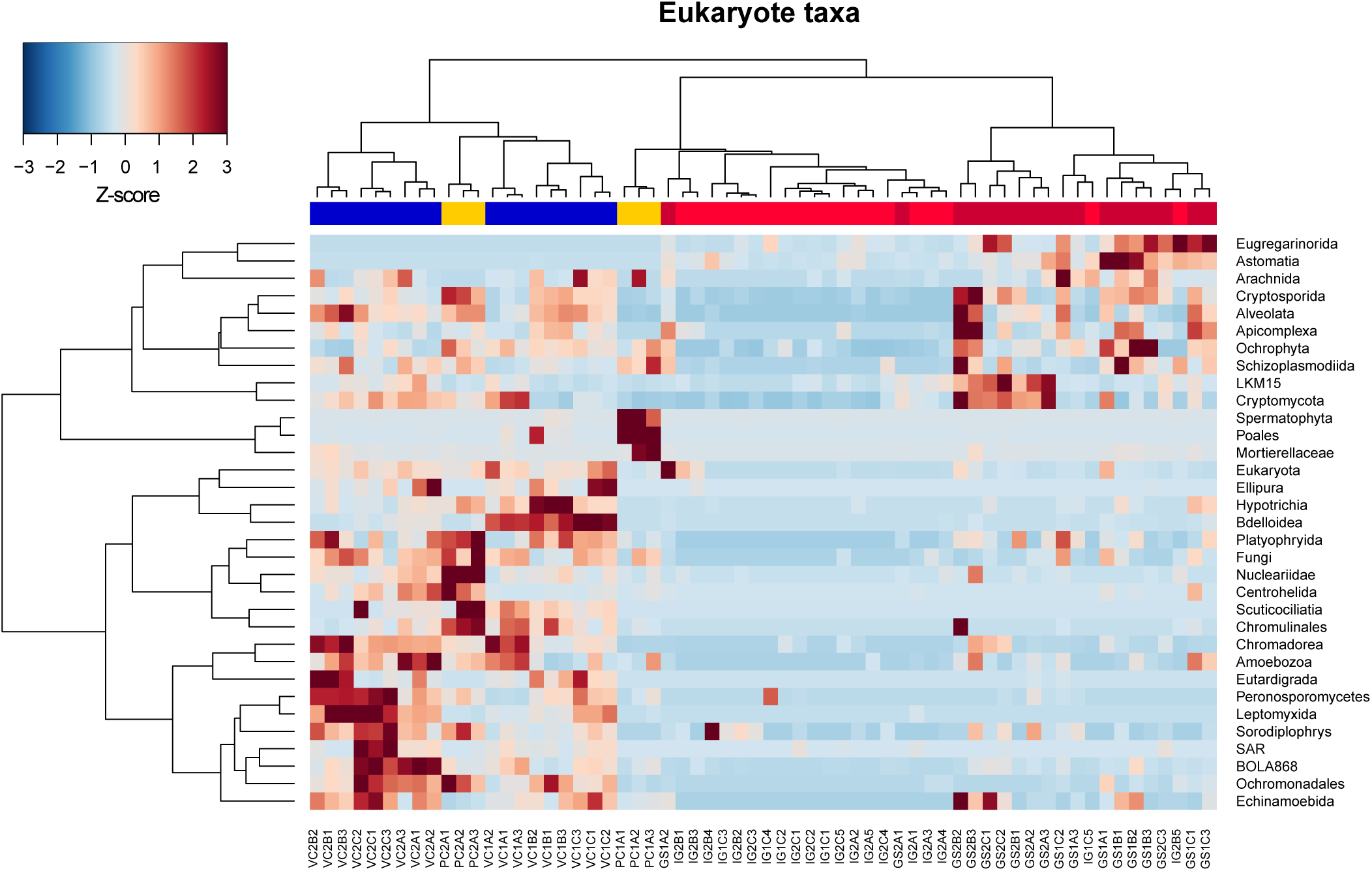
Binned taxa filtered for those achieving 1% abundance in 5% of samples in the rarefied ASV table plotted by Ward’s clustering of scaled abundance shows a distinct division in relative enrichment of microeukaryotes. Compost samples PC and VC (yellow and blue respectively) are generally enriched across a larger number of taxonomic bins relative to GS samples (red).

The Chromadorea and Ellipura percent relative abundance in GS samples were approximately 1.88% and 0.02% of rarefied reads respectively, while significantly greater proportions of these two organisms were found in VC samples (castings) (Wilcoxon: p<0001). *Mortierellaceae* were most prevalent in PC (7.55%) while <1% in VC and GS, and *Thraustochytriaceae* were <1% for all sample types (Table S4).

Nematodes contain all desaturase and elongase enzymes to denovo produce up to 20C length FAs, including 20:5ω3 EPA and 20:4n6 ARA, which were the uniquely enriched PUFAs found among our GS samples (61). Therefore, we cannot exclude that the prevalence of nematodes passing through the earthworm intestinal tract and expelled with the castings may explain the detection of PUFAs in the GS. However, since PC samples also contain a smaller but detectable amount of nematode 18S rRNA signal, and yet are devoid of any detectable PUFA, then it is unclear whether the nematodes in the earthworm composting system where we sampled are sequestering PUFA. Previous studies have shown that *Caenorhabditis elegans* may contain anywhere from 1% to 20% of total lipid PUFA, depending on growth temperature (63). These are largely due to an abundance of 18C FAs, namely a signature peak at 18:1ω7 (*cis*-vaccenic acid) in all lipid fractions – phosphatidylcholine, phatidylethanolamine – but to a lesser degree also EPA and ARA (59, 61, 63, 64). However, none of our sample peaks were annotated for 18:1ω7, which would be expected if nematode lipids (or otherwise aerobic bacteria) were contributing to the pool of measured fatty acids in our samples. Furthermore, earthworms respond to nematode infestations that occur in the coelomic cavity and near the nephridia by encapsulating the tiny worms in cysts called “brown bodies” and destroying them using reactive oxygen species (57). Alternatively, the earthworm itself may be able to produce PUFAs, as recent genomic analysis has revealed genes encoding “methyl”-end desaturase homologs to the Δ12 (ω6) and Δ15 (ω3) desaturases in Oligochaeta (18). Little is known about the ability of invertebrates to produce long-chain PUFA, especially the amount and rate that this might occur. Evidence from genomic analysis attests to wide-spread occurrence of elongation and desaturation enzymes, but only a handful of earlier publications have indicated some limited de novo production such as in Collembola (springtails), *Daphnia* (water fleas) and nematodes (20, 59, 61). Still, Arthropoda exhibit poor fecundity and decline in fitness when their dietary supply of pre-formed PUFA are absent or restricted, despite encoding the competent enzymes for de novo production (20, 59). In earthworms, the increasing concentration of PUFAs from bulk soil to gut soil, and finally peaking in muscle tissue, which was interpreted as trophic transfer from soil microorganisms to the earthworm (28), may well work in the opposite direction. The earthworm may be producing PUFA within its own tissue, perhaps as precursors to eicosanoids or for cellular membrane permeability, and as the epithelial cells shed into the gut, then PUFAs appear to concentrate in gut soil. However, in this prior study the entire unbroken intestinal track was surgically isolated from body, and it challenges any notion that somatic cell metabolites should be leaked into the gut soil, and rather strengthens the original interpretation. To definitively rule out earthworm-derived PUFAs, further work should isolate metabolic activity using approaches such as stable isotope probing with heavy isotopes of carbon and hydrogen.

### Conclusions

In sum, we found that the earthworm gut soil ecosystem is distinct from the input compost and output castings according to the type and abundance of organisms present, as well as in the fatty acid metabolite profiles. Amplicon sequencing of the KS functional domain on the *pfaA* gene as part of the PUFA synthase complex illuminated the sparse and phylogenetically narrow distribution of putative PUFA producing bacterial taxa, especially the ASVs from GS samples, which all unambiguously belonged to strains of *Shewanella*. Most surprising was that while the 20C PUFAs were found among the total lipid pool of the GS samples and absent in both compost sample types, the PfaA-KS sequence abundance pattern was exactly the contrary. Samples in which the largest number and diversity of PfaA-KS sequences were recovered were VC, which are the castings or feces of the earthworms. These samples contained taxa also found among the GS samples which were implicated as PUFA-competent based on PfaA-KS presence, but VC samples also contained other PfaA-KS-positive taxa known to be wide-spread free-living organisms. Based on 16S and 18S rRNA taxonomic profiling, the GS samples diverged from the compost samples based on presence and abundance of ASVs, suggestive of a unique ecosystem that is maintained by the conditions of the earthworm gut. Eukaryotic origin of the PUFAs is possible but not well supported, since organisms such as nematodes and collembola failed to show other signature 18C fatty acid biomarkers, and were not well-represented among the amplicon sequence pool, especially of the GS samples. The overall picture appears to be that earthworms may aggregate the ARA and EPA PUFAs stemming from bacterial metabolism, or they produce these lipids from their own genomically-encoded enzymes. Conditions in the intestinal tract may activate PUFA-producing taxa like *Shewanella* that proliferate in the earthworm gut ecosystem during passage of environmental soil and bacteria.

## Material and Methods

### Materials

| Method | Reagent |
| --- | --- |
| Lipid extraction | - Chloroform<br>- Methanol<br>- dH<sub>2</sub>O<br>- Sodium chloride (NaCl) |
| Basic derivatization | - Hexane<br>- Potassium hydroxide (KOH)<br>- Methanol<br>- Sodium Bisulphate (NaHSO<sub>4</sub>) |
| Acid derivatization | - Sulfuric acid (H<sub>2</sub>SO<sub>4</sub>)<br>- Methanol<br>- Hexane<br>- Sodium bicarbonate (NaHCO<sub>3</sub>)<br>- Sodium sulphate anhydrous (Na<sub>2</sub>SO<sub>4</sub>) |
| Nucleic acid extraction:<br>ZymoBIOMICS™ MagBead DNA/RNA Kit | - ZymoBIOMICS™ MagBinding Beads<br>- DNA/RNA Shield™<br>- DNA/RNA Lysis Buffer<br>- DNA/RNA Prep Buffer<br>- MagBead DNA/RNA Wash 1<br>- MagBead DNA/RNA Wash 2<br>- DNase/RNase-Free Water<br>- DNase I 3 (lyophilized) 250 U<br>- DNA Digestion Buffer 4 ml 4 ml<br>- Proteinase K 3 (lyophilized) & Storage Buffer |

### Equipment

| Equipment | Company |
| --- | --- |
| CentriVap Benchtop Vacuum concentrator | Labconco |
| HP-5ms Ultra Inert Column | Aglient |
| 7890A GC system | Aglient |
| Fastprep-96 | MP |
| MiSeq System | Illumina |

### Experimental design and sample collection

Adult *Eisenia fetida* were collected from Vermigrand GmbH in Absdorf, Austria together with different compost samples, designated Pre-Compost and Vermicompost. Pre-Compost (PC) is compost consisting of horse manure and thermophilic compost, it is laid freshly onto the existing compost every week to feed the worms. Vermicompost (VC) is worm castings and is used as an organic fertilizer. Samples were collected on 3 specific days within a one-week period, for 2 separate weeks. This sampling scheme was chosen to capture the possible effects within a week of composting, since new Pre-compost is laid onto existing soil once a week.

Collected worms were kept in their natural soil composition until the dissection, taking place not more than two hours later. For dissection, the worms were put into a petri dish and cleaned from residual soil before being euthanized in hot water (70°C) for 20s. Worms were then dissected aseptically under a stereomicroscope to recover soil from the intestinal tract (called “Gut soil” or GS). Gut soil from ten to fifteen worms was collected to recover at least 275 mg wet gut soil per replicate. To collect castings of *E. fetida*, worms were cleaned externally with deionized water and then separated into individual culture boxes with ventilation for 48 hours. The castings produced the worms were collected, weighed, and then pooled to recover at least 275 mg wet weight. Samples were frozen immediately after collection, freeze dried on the next day, and stored at –20°C until further use.

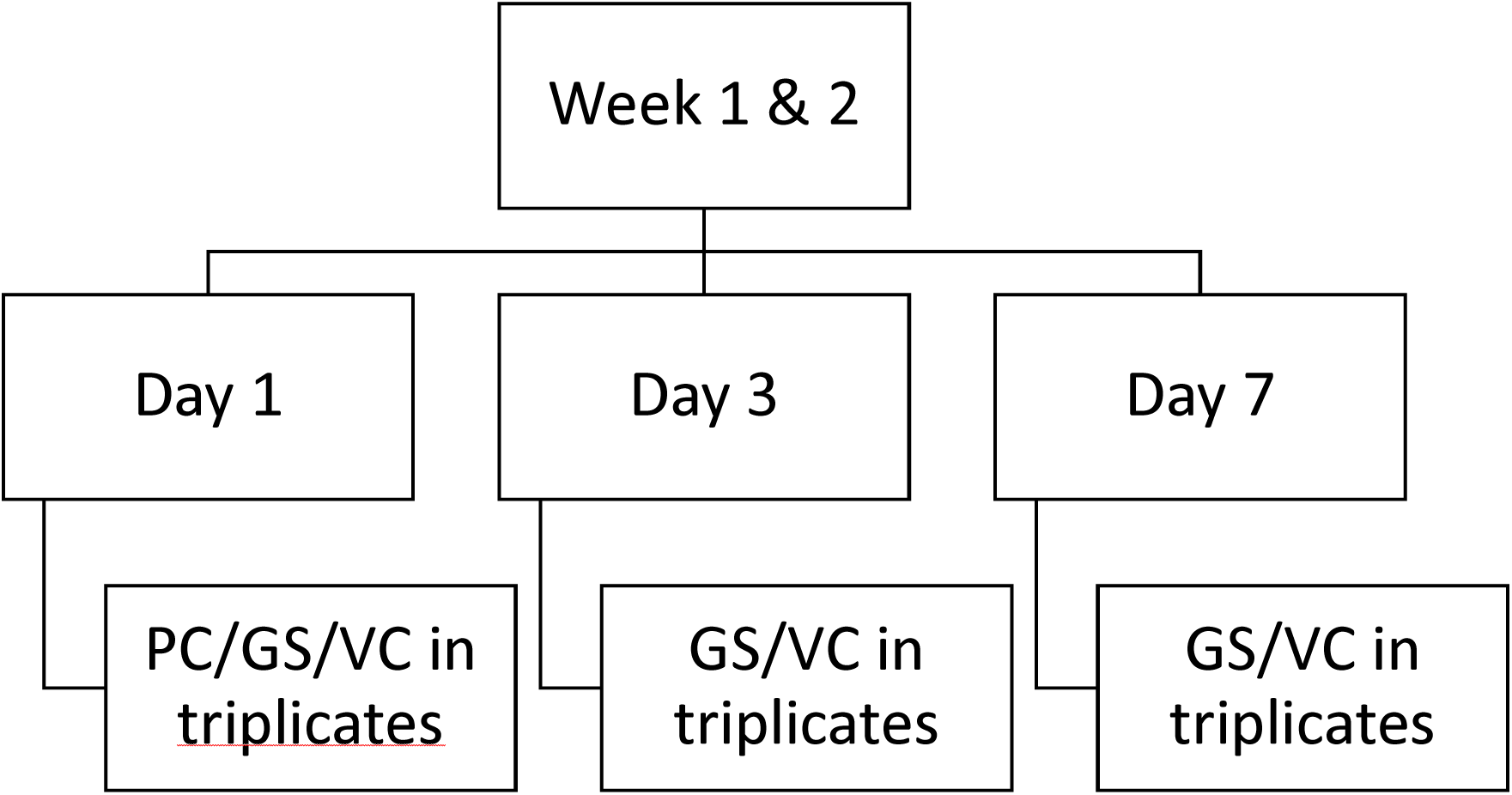

Every sample was given a unique identifier for tracking the sample during the analysis as shown below:

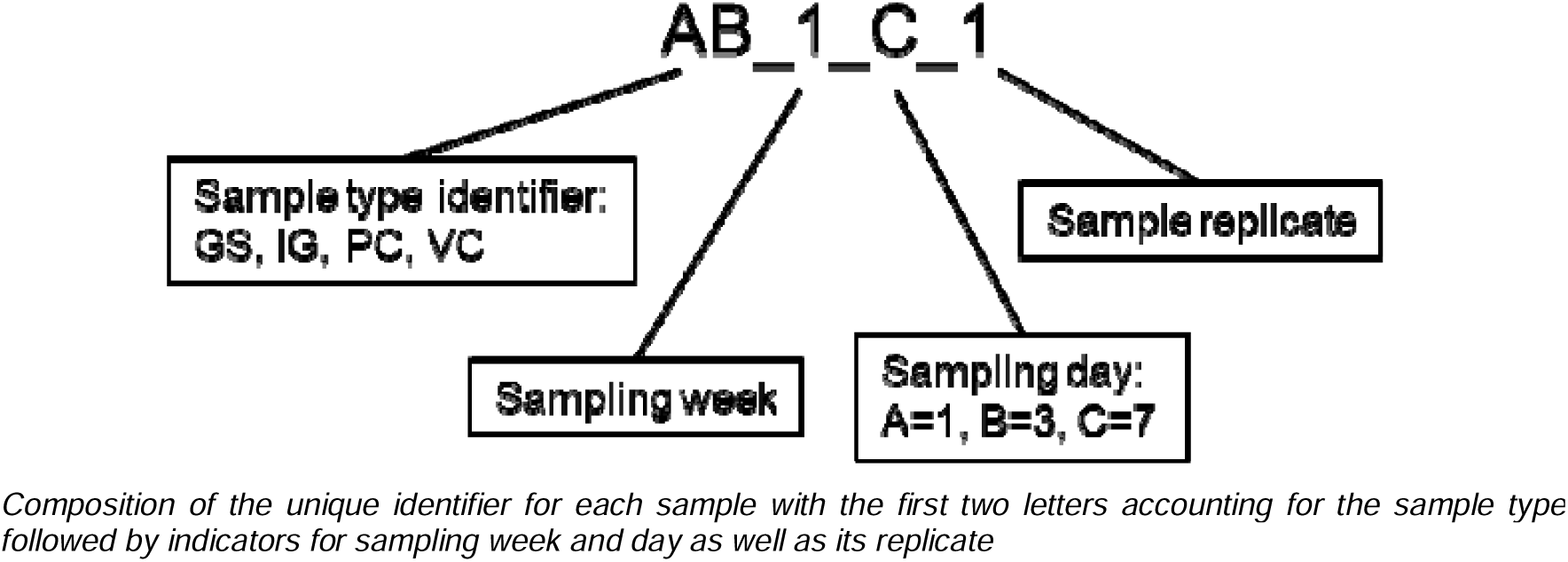

### Lipid extraction

The Folch extraction method was carried out following the procedure described by Folch et al. (65): 70 mg dry sample (PC, VC and GS) was extracted with 1.4_mL of chloroform:methanol (2:1) vortexing for two_minutes. The mixture was centrifuged at 4000_rpm for 10_min before collecting the supernatant. The extraction process was carried out three times on the same sample and pooled. The collected organic layers were purified washing with 1 mL dH_2_O and a spatula tip of NaCl. The mixture was vortexed for one minute and centrifuged at 4000_rpm for 10_minutes. The chloroform layer that contains the extracted lipids was recovered and evaporated in a Labconco CentriVap Benchtop Vacuum concentrator under reduced pressure at 10_°C. The lipid content was determined gravimetrically and calculated as weight percentage of dry biomass. Lipid extracts obtained were directly used for derivatization.

#### Basic derivatization

Basic derivatization was carried out by resuspending 10 mg lipid extracts in 100 µL of hexane. After vortexing, to completely dissolve the extract, 50 µL of 2 M KOH solution in methanol was added and vortexed for one minute. The sample was incubated for five minutes at room temperature. 125 mg of sodium bisulphate was added to the sample, vortexed and centrifuged at 4000 rpm for five minutes. 100 µL of supernatant was collected, mixed with 400 µL of hexane and filtered through a 0.22 µm PTFE filter. The filter was washed with an additional 100 µL of Hexane.

#### GC-MS method

FAMEs were separated using a HP-5ms Ultra Inert Column (30 m x 250 μm x 0.25 μm) (Agilent, CA, USA). 1_μL sample was injected in spitless mode and an injector temperature of 280 °C. The initial oven temperature was set at 150_°C for 1_min and the temperature was gradually raised to 220_°C at 3_°C/min with a final increase to 300 °C for 3 min. Helium was used as carrier gas at a constant column flow rate of 2.52_mL/min. The GC to MS interface temperature was fixed at 280 °C and an electron ionization system was set on the MS in scan mode. The mass range evaluated was 50–600 m/z, where MS quad and source temperatures were maintained at 150 °C and 230 °C respectively. To search and identify each fatty acid, the NIST-MS Library (2.2) was used. It was also used to measure the relative percentage of each compound and relative peak areas of the total ionic chromatogram were used. Additionally, a commercial standard (Supelco 37 Component FAME Mix cat.#47885) was used to identify peaks of the fatty acids by comparing retention times to those of known fatty acids in the commercial standard.

#### Statistical analysis of identified fatty acids

All extractions were performed in biological triplicate for each time point within a week. The results were expressed as mean ± standard deviation while the effect of the extraction method and the extraction conditions on lipid recovery was analyzed by using two-way ANOVA followed by Tukey post-hoc test (statistically significance at p < 0.05). Statistical analysis was performed with GraphPad Prism 8 (8.0.1) (GraphPad Software, San Diego, CA, USA). To test differences between the two sampling weeks and to discriminate differences in fatty acid composition between samples, the data were transformed by using center-log ratio and then evaluated using a Kolmogorov-Smirnov t-test and Euclidean distance-based ordination respectively. Between-group differences were assessed by permutational ANOVA by explaining community distances based by sampling week interacting with sampling day and sample type: (Response = adonis2 (community data ∼ sampling week * sampling day * sample type)). Every analysis was performed with R version 4.1.1 using the “vegan” package (v2.6-4) (66).

### Nucleic acids extraction and sequencing

DNA and RNA extraction was carried out by using the ZymoBIOMICS^TM^ MagBead DNA/RNA Kit (67). Samples were weighed at roughly equivalent starting biomass (approximately 20 mg dry weight when sample quantity allowed) into prefilled tube strips (8 x 12) containing 750 µl DNA/RNA Shield^TM^ (ZymoBIOMICS) and 40-400 µm glass beads (Macherey-Nagel) and arranged on a 96-well rack. Homogenization was conducted on an MP Fastprep-96 with one minute on maximum speed and five minutes rest, which was repeated five times. DNA and RNA were purified from the same input material following the kit protocol, yielding two separate eluates for DNA and RNA respectively. The concentrations were measured with the Qubit 1x dsDNA HS Assay Kit and RNA HS Assay Kit. The detailed protocol for amplicon generation and sequencing are published by Pjevac et al. (68). In brief, a two-step PCR was performed for amplicon generation and to add linkers as well as barcodes. Amplicons were generated for the V4 region of the 16S (515F: GTG YCA GCM GCC GCG GTA A; 806R: GGA CTA CNV GGG TWT CTA AT) and 18S (TAReuk454FWD1: CCA GCA SCY GCG GTA ATT CC; TAR eukREV3mod: ACT TTC GTT CTT GAT YRA TGA) rRNA genes as well as the PfaA-KS domain region of the *pfa* gene (PfaA-KS-Fw: TGG GAA GAR AWT TCC C; PfaA-KS-Rv: GTR CCN GTR CNG CTT C), part of the Pfa-Synthase complex (33). Linker sequences from primer to barcode were forward GCT ATG CGC GAG CTG C and reverse TAG CGC ACA CCT GGT A. Cycling conditions for the 16S and 18S rRNA amplicons were: initial denature 94° C for 3 mins, and then denaturing at 95° C for 30 s, annealing at 55° C for 30 s, and extension at 72° C for 60 s. Cycling conditions for the PfaA-KS amplicons were: initial denature at 94° C for 3 min, and then denaturing at 94° C for 45 s, annealing at 50.3° C for 30 s, and extension at 72° C for 90 s. The first PCR for all-three primer sets was done with 30 cycles and 0,25 µM primer concentration while the second PCR was done with 7 cycles for the 16S and 18S rRNA amplicons and 15 cycles for the PfaA-KS amplicons using 0,8 µM primer concentration. Sequencing was performed by the Joint Microbiome Facility (JMF) on an Illumina MiSeq using 2×300 base pair sequencing (69).

### Analysis of amplicon sequencing data

#### In-house generated data

Input data were filtered for PhiX contamination with BBDuk (70). Demultiplexing was performed with the python package demultiplex allowing 1 mismatch for barcodes and 2 mismatches for linkers (71). Primers were verified with the python package demultiplex allowing 2 and 2 mismatches for forward and reverse primers, respectively. Barcodes, linkers, and primers were trimmed off using BBDuk with 47 and 48 bases being left-trimmed for F.1/R.2 and F.2/R.1, respectively allowing a minimum length of 220 base pairs. Around 80% of reads across all samples passed all trimming, filtering and denoising criteria.

#### Amplicon Sequence Variants (ASV) determination

ASVs for 16S and 18S sequences were generated by in-house sequencing facility using a standard workflow with DADA2 (72). The PfaA-KS sequences were handled separately according to the following steps: (i) Forward and reverse orientated sequences were kept separated based on primer (F.1/F.2 and R.1/R.2), and merged using PEAR (73) with minimum overlap (-o) of 5, minimum trim length (-t) of 300, maximum assembly length (-m) of 600, minimum quality score (-q) of 30, and maximum uncalled bases allowed (-u) of 0; (ii) The forward-orientated FASTQ reads starting with the reverse primer (R: merged from R1/R2) were reverse complemented, and then all merged sequences were concatenated per sample and used for ASV calling DADA2; (iii) Filtering was conducted based on quality profile plots: length truncated to 475 nt, maxN=0, maxEE=5, truncQ=3, rm.phix=TRUE. After merging and filtering, 11 samples were discarded (01, 16, 24, 27, 40, 41, 42, 50, 52, 63, and 64) because no reads passed filtering; (iv) The remaining 54 samples were used for error modelling and ASV calling, yielding 66 unique biological PfaA-KS sequence ASVs. The sample ASV table and ASV sequences were exported for further analyses.

#### Taxonomy assignments

SSU rRNA ASVs were classified using SINA version 1.6.1 and the SILVA database SSU Ref NR 99 release 138.1 (74, 75). The PfaA-KS sequences were classified using BLASTn and BLASTx against the nucleotide (nt) and non-redundant protein (nr) databases respectively and the top hit extracted for putative assignment (Table S5). The results from BLASTn were primarily used unless the identity and query coverage fell below 80%, in which the BLASTx results were consulted. Sixteen samples remained with ambiguous assignments (“uncultured organism”) and for these the BLASTx (nr) search was narrowed to Bacterial references only, which resolved the assignments, with a minimum percent identity of 81% and full query coverage (see Tables S6 & S7 for all BLAST output). Top-hits for each amplicon variant were used for the final assignment, and the full taxonomy lineage was retrieved from the Genome Taxonomy Database, release 202 metadata (76–79).

#### Alignment and tree-building

The PfaA-KS sequence variants from the original study (33) were accessed from the NCBI GenBank PopSet 303162115 (446 total entries) and reannotated following the same procedures with BLASTn and BLASTx as for our in-house generated PfaA-KS ASVs (Table S8). These sequences were combined with our in-house generated PfaA-KS ASVs. The *Escherichia coli fabF* gene was used as an outgroup to root the phylogeny. Sequences were aligned with MAFFT (v7.520) using the –-localpair option and –-maxiterate 1000 (80, 81). Phylogenetic inference was made using IQ-TREE (multicore version 1.6.12) (82–85), first using model selection, and then using the best-fit model with the following parameters: –m TIM2e+R5 –bb 1000 –nt 4 –redo. The resulting tree was visualized using iTOL (86).

#### Genomic mining of PfaA-KS primer matches

To validate the use of universal primers designed for marine bacteria using a region on the ketoacyl-synthase domain of the *pfaA* gene (Shulse & Allen 2010), all available whole genomes of archaea and bacteria were downloaded from the NCBI GenBank archive. Putative PUFA synthase regions were found using a sequence-naïve approach by searching for tandem repeats of the phosphopantetheine attachment site for the acyl-carrier-protein domain. The genomic region containing the full putative PfaA-KS domain was extracted from reference genomes of positive hits. The blastn command line tool was used to map all forward and reverse sequence possibilities of the degenerate primers, setting word-size to 7 and qcov_hsp_perc (percent-query-coverage per high scoring pair) to 50 to accommodate loose configurations of primer matches. The matches were annotated as to whether or not the last three bases on the 3’ end aligned to the reference, and the out6 formats of all hits compiled into a table, one entry per reference, with the final header: qseqid, sseqid, length, mismatch, frames, qstart, qend, sstart, send, sseq, and the custom column for 3’-codon-matches for forward and reverse respectively. The originally extracted putative PfaA-KS domains from the reference genomes were aligned using MAFFT and then phylogenetic inference made using IQ-TREE, with the resulting tree file exported and visualized in iTOL. The entries were colored according to the taxonomic lineage. The leaves were annotated as to their primer pair match, 3’-end coverage, and the predicted insert size to visualize amplicons that could be theoretically obtained from across the entire phylogenetic space.

#### Analysis and statistics

All analysis on the ASV abundance tables were handled in R version 4.1.1 including the following packages: {decontam} (v1.14.0) was used for removing contaminants (87); sampling depth was normalised with {DESeq2} (v1.36.0) using a variance stabilizing transformation (88); compositional transformations using center-log-ratio (CLR) and estimation of effect sizes were handled with {propr}, {vegan}, and {ALDEx2} (53, 89–92). An effect size >1 was considered as explanatory. The base R {stats} was used for clustering samples as well as pair-wise comparisons with the Wilcoxon test. Additionally, to reduce the rate of type-I errors, multiple testing corrections were done with the Benjamini-Hochberg method. False discovery rate (FDR)_<_0.05 was considered as statistically significant (93). {phyloseq} (v1.40.0) was used for alpha– and beta-diversity calculation while rarefaction for alpha diversity calculation was done with {vegan} (2.6-4) (89, 94). Heatplots were generated with {made4} (v1.68.0) (95), {massageR} and {gplots} (96, 97); dataframe transformation and reformatting were handled with {reshape2}, {dplyr} (v1.0.10), {tidyr} (v1.2.1) (98, 99), and plotting conducted with {ggplot2} using {wesanderson} color palettes (100, 101).

#### Additional Information

Sequence reads for 16S, 18S, and PfaA-KS amplicons have been uploaded to NCBI SRA under BioProject ID PRJNA1152521

