## Supplemental Figures 1-10 for "*Shewanella* is a putative producer of polyunsaturated fatty acids in the gut soil of the composting earthworm *Eisenia fetida*"

**Figure S1.** Amplicon rarefaction curves for rRNA sequences

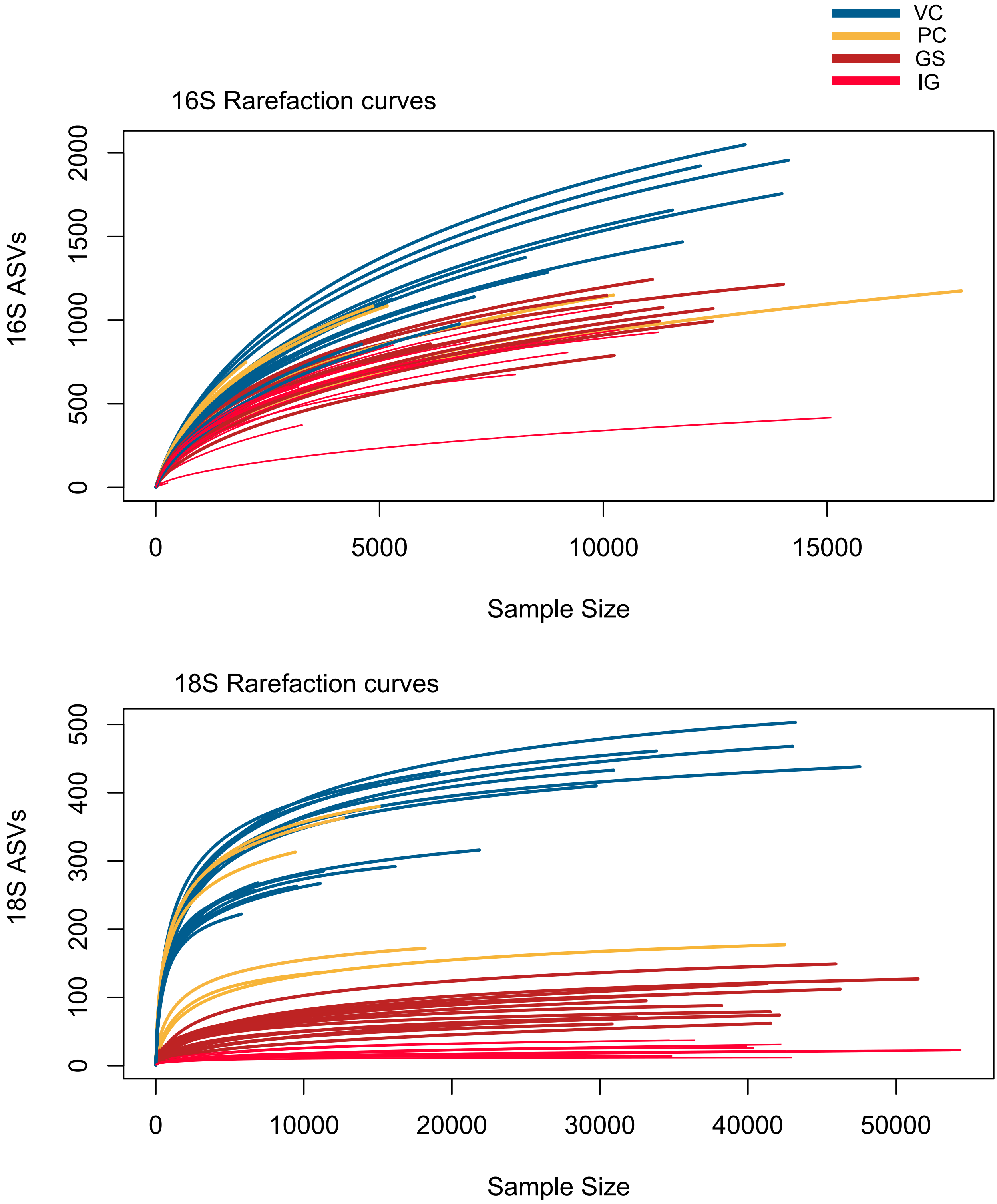

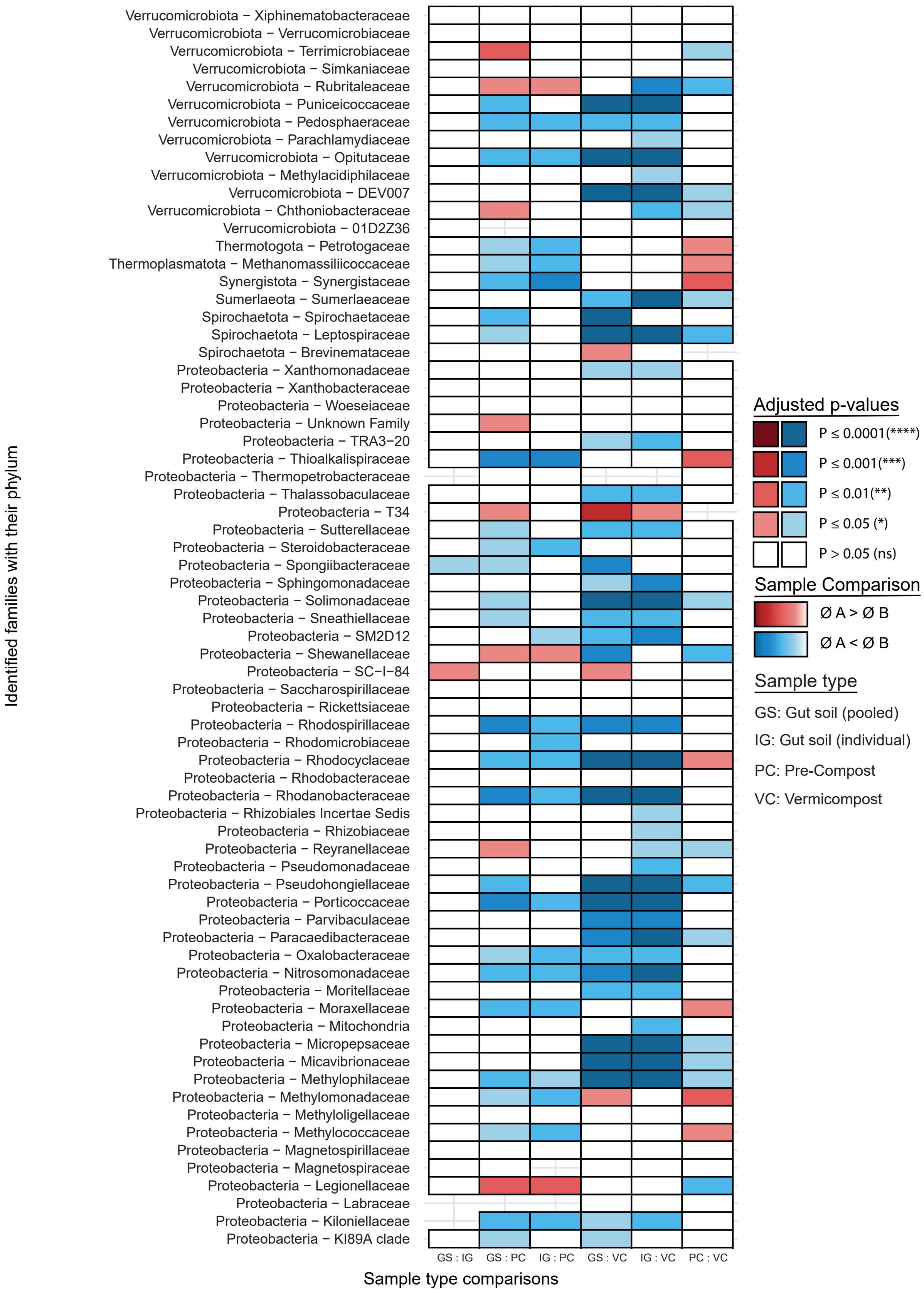

Verrucomicrobiota − Xiphinematobacteraceae
Verrucomicrobiota − Verrucomicrobiaceae
Verrucomicrobiota − Terrimicrobiaceae
Verrucomicrobiota − Simkaniaceae
Verrucomicrobiota − Rubritaleaceae
Verrucomicrobiota − Puniceicoccaceae
Verrucomicrobiota − Pedosphaeraceae
Verrucomicrobiota − Parachlamydiaceae
Verrucomicrobiota − Opitutaceae
Verrucomicrobiota − Methylacidiphilaceae
Verrucomicrobiota − DEV007
Verrucomicrobiota − Chthoniobacteraceae
Verrucomicrobiota − 01D2Z36
Thermotogota − Petrotogaceae
Thermoplasmatota − Methanomassiliicoccaceae
Synergistota − Synergistaceae
Sumerlaeota − Sumerlaeaceae
Spirochaetota − Spirochaetaceae
Spirochaetota − Leptospiraceae
Spirochaetota − Brevinemataceae
Proteobacteria − Xanthomonadaceae
Proteobacteria − Xanthobacteraceae
Proteobacteria − Woeseiaceae
Proteobacteria − Unknown Family
Proteobacteria − TRA3−20
Proteobacteria − Thioalkalispiraceae
Proteobacteria − Thermopetrobacteraceae
Proteobacteria − Thalassobaculaceae
Proteobacteria − T34
Proteobacteria − Sutterellaceae
Proteobacteria − Steroidobacteraceae
Proteobacteria − Spongiibacteraceae
Proteobacteria − Sphingomonadaceae
Proteobacteria − Solimonadaceae
Proteobacteria − Sneathiellaceae
Proteobacteria − SM2D12
Proteobacteria − Shewanellaceae
Proteobacteria − SC−I−84
Proteobacteria − Saccharospirillaceae
Proteobacteria − Rickettsiaceae
Proteobacteria − Rhodospirillaceae
Proteobacteria − Rhodomicrobiaceae
Proteobacteria − Rhodocyclaceae
Proteobacteria − Rhodobacteraceae
Proteobacteria − Rhodanobacteraceae
Proteobacteria − Rhizobiales Incertae Sedis
Proteobacteria − Rhizobiaceae
Proteobacteria − Reyranellaceae
Proteobacteria − Pseudomonadaceae
Proteobacteria − Pseudohongiellaceae
Proteobacteria − Porticoccaceae
Proteobacteria − Parvibaculaceae
Proteobacteria − Paracaedibacteraceae
Proteobacteria − Oxalobacteraceae
Proteobacteria − Nitrosomonadaceae
Proteobacteria − Moritellaceae
Proteobacteria − Moraxellaceae
Proteobacteria − Mitochondria
Proteobacteria − Micropepsaceae
Proteobacteria − Micavibrionaceae
Proteobacteria − Methylophilaceae
Proteobacteria − Methylomonadaceae
Proteobacteria − Methyloligellaceae
Proteobacteria − Methylococcaceae
Proteobacteria − Magnetospirillaceae
Proteobacteria − Magnetospiraceae
Proteobacteria − Legionellaceae
Proteobacteria − Labraceae
Proteobacteria − Kiloniellaceae
Proteobacteria − KI89A clade
Identified families with their phylum
GS : IG
GS : PC
IG : PC
GS : VC
IG : VC
PC : VC
Sample type comparisons
Adjusted p-values
P ≤ 0.0001(****)
P ≤ 0.001(***)
P ≤ 0.01(**)
P ≤ 0.05 (*)
P > 0.05 (ns)
Sample Comparison
Ø A > Ø B
Ø A < Ø B
Sample type
GS: Gut soil (pooled)
IG: Gut soil (individual)
PC: Pre-Compost
VC: Vermicompost

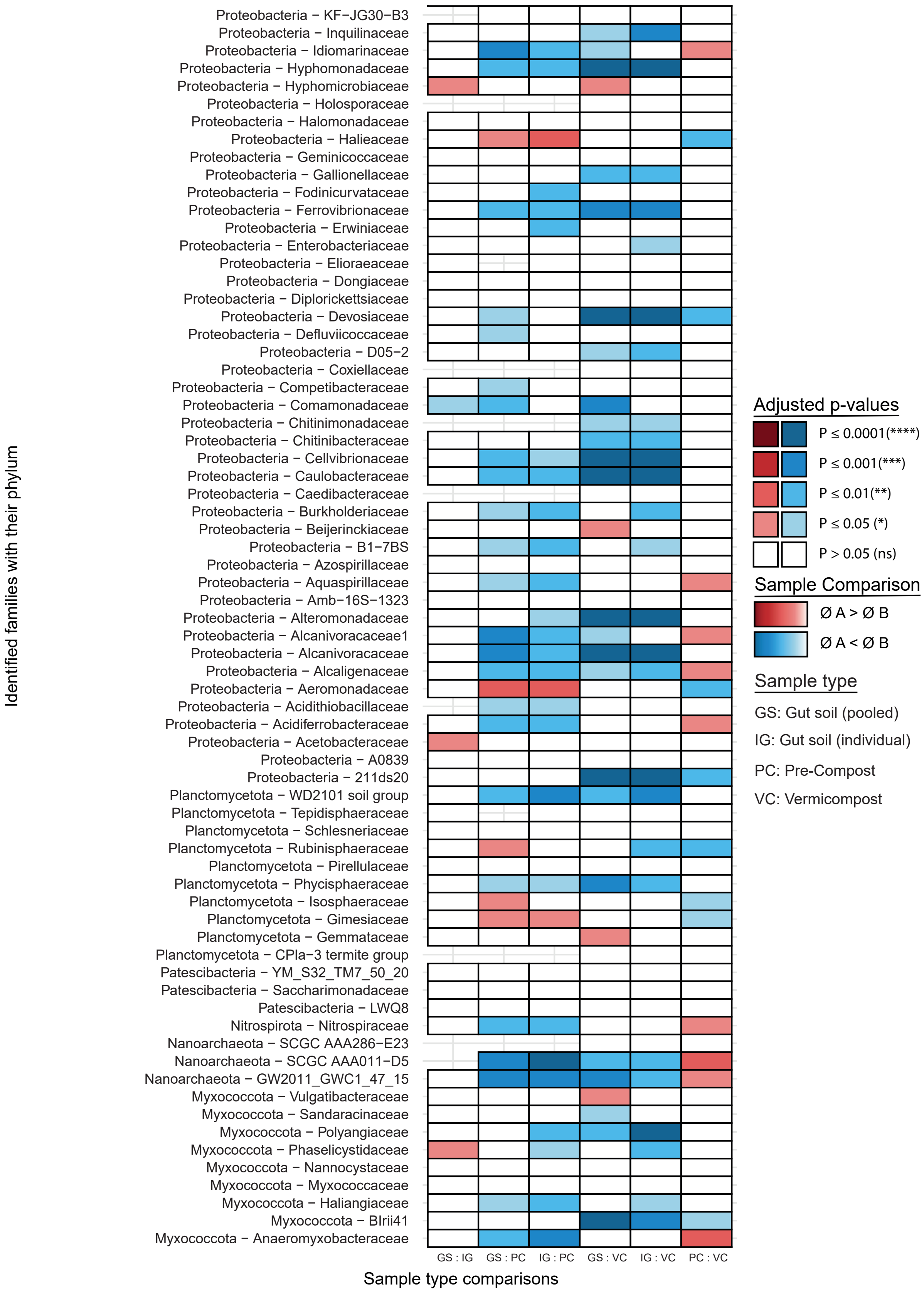
Identified families with their phylum
Proteobacteria − KF−JG30−B3
Proteobacteria − Inquilinaceae
Proteobacteria − Idiomarinaceae
Proteobacteria − Hyphomonadaceae
Proteobacteria − Hyphomicrobiaceae
Proteobacteria − Holosporaceae
Proteobacteria − Halomonadaceae
Proteobacteria − Halieaceae
Proteobacteria − Geminicoccaceae
Proteobacteria − Gallionellaceae
Proteobacteria − Fodinicurvataceae
Proteobacteria − Ferrovibrionaceae
Proteobacteria − Erwiniaceae
Proteobacteria − Enterobacteriaceae
Proteobacteria − Elioraeaceae
Proteobacteria − Dongiaceae
Proteobacteria − Diplorickettsiaceae
Proteobacteria − Devosiaceae
Proteobacteria − Defluviicoccaceae
Proteobacteria − D05−2
Proteobacteria − Coxiellaceae
Proteobacteria − Competibacteraceae
Proteobacteria − Comamonadaceae
Proteobacteria − Chitinimonadaceae
Proteobacteria − Chitinibacteraceae
Proteobacteria − Cellvibrionaceae
Proteobacteria − Caulobacteraceae
Proteobacteria − Caedibacteraceae
Proteobacteria − Burkholderiaceae
Proteobacteria − Beijerinckiaceae
Proteobacteria − B1−7BS
Proteobacteria − Azospirillaceae
Proteobacteria − Aquaspirillaceae
Proteobacteria − Amb−16S−1323
Proteobacteria − Alteromonadaceae
Proteobacteria − Alcanivoracaceae1
Proteobacteria − Alcanivoracaceae
Proteobacteria − Alcaligenaceae
Proteobacteria − Aeromonadaceae
Proteobacteria − Acidithiobacillaceae
Proteobacteria − Acidiferrobacteraceae
Proteobacteria − Acetobacteraceae
Proteobacteria − A0839
Proteobacteria − 211ds20
Planctomycetota − WD2101 soil group
Planctomycetota − Tepidisphaeraceae
Planctomycetota − Schlesneriaceae
Planctomycetota − Rubinisphaeraceae
Planctomycetota − Pirellulaceae
Planctomycetota − Phycisphaeraceae
Planctomycetota − Isosphaeraceae
Planctomycetota − Gimesiaceae
Planctomycetota − Gemmataceae
Planctomycetota − CPla−3 termite group
Patescibacteria − YM_S32_TM7_50_20
Patescibacteria − Saccharimonadaceae
Patescibacteria − LWQ8
Nitrospirota − Nitrospiraceae
Nanoarchaeota − SCGC AAA286−E23
Nanoarchaeota − SCGC AAA011−D5
Nanoarchaeota − GW2011_GWC1_47_15
Myxococcota − Vulgatibacteraceae
Myxococcota − Sandaracinaceae
Myxococcota − Polyangiaceae
Myxococcota − Phaselicystidaceae
Myxococcota − Nannocystaceae
Myxococcota − Myxococcaceae
Myxococcota − Haliangiaceae
Myxococcota − BIrii41
Myxococcota − Anaeromyxobacteraceae
GS : IG
GS : PC
IG : PC
GS : VC
IG : VC
PC : VC
Sample type comparisons
Adjusted p-values
P ≤ 0.0001(****)
P ≤ 0.001(***)
P ≤ 0.01(**)
P ≤ 0.05 (*)
P > 0.05 (ns)
Sample Comparison
Ø A > Ø B
Ø A < Ø B
Sample type
GS: Gut soil (pooled)
IG: Gut soil (individual)
PC: Pre-Compost
VC: Vermicompost

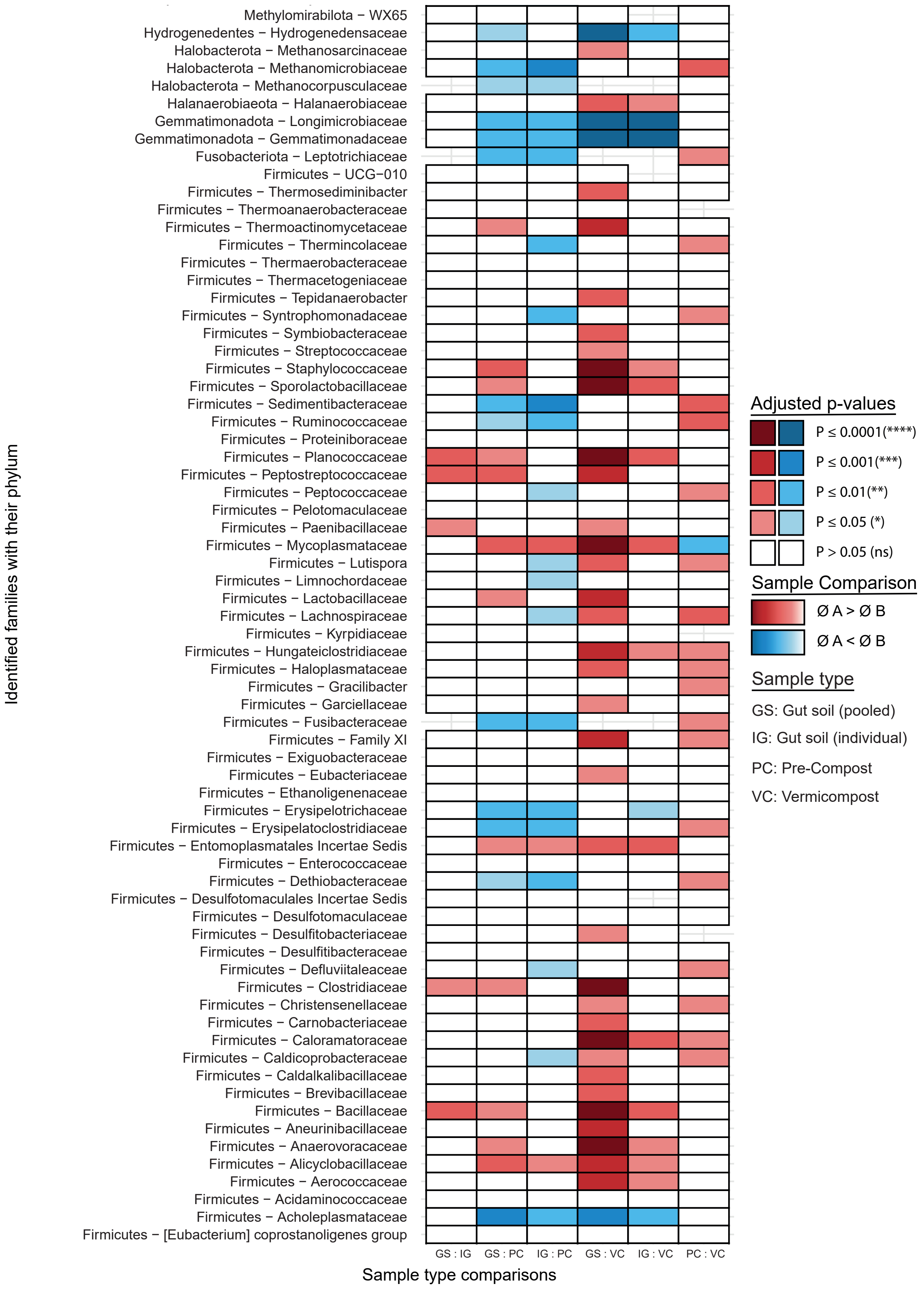

Identified families with their phylum
Methylomirabilota − WX65
Hydrogenedentes − Hydrogenedensaceae
Halobacterota − Methanosarcinaceae
Halobacterota − Methanomicrobiaceae
Halobacterota − Methanocorpusculaceae
Halanaerobiaeota − Halanaerobiaceae
Gemmatimonadota − Longimicrobiaceae
Gemmatimonadota − Gemmatimonadaceae
Fusobacteriota − Leptotrichiaceae
Firmicutes − UCG−010
Firmicutes − Thermosediminibacter
Firmicutes − Thermoanaerobacteraceae
Firmicutes − Thermoactinomycetaceae
Firmicutes − Thermincolaceae
Firmicutes − Thermaerobacteraceae
Firmicutes − Thermacetogeniaceae
Firmicutes − Tepidanaerobacter
Firmicutes − Syntrophomonadaceae
Firmicutes − Symbiobacteraceae
Firmicutes − Streptococcaceae
Firmicutes − Staphylococcaceae
Firmicutes − Sporolactobacillaceae
Firmicutes − Sedimentibacteraceae
Firmicutes − Ruminococcaceae
Firmicutes − Proteiniboraceae
Firmicutes − Planococcaceae
Firmicutes − Peptostreptococcaceae
Firmicutes − Peptococcaceae
Firmicutes − Pelotomaculaceae
Firmicutes − Paenibacillaceae
Firmicutes − Mycoplasmataceae
Firmicutes − Lutispora
Firmicutes − Limnochordaceae
Firmicutes − Lactobacillaceae
Firmicutes − Lachnospiraceae
Firmicutes − Kyrpidiaceae
Firmicutes − Hungateiclostridiaceae
Firmicutes − Haloplasmataceae
Firmicutes − Gracilibacter
Firmicutes − Garciellaceae
Firmicutes − Fusibacteraceae
Firmicutes − Family XI
Firmicutes − Exiguobacteraceae
Firmicutes − Eubacteriaceae
Firmicutes − Ethanoligenenaceae
Firmicutes − Erysipelotrichaceae
Firmicutes − Erysipelatoclostridiaceae
Firmicutes − Entomoplasmatales Incertae Sedis
Firmicutes − Enterococcaceae
Firmicutes − Dethiobacteraceae
Firmicutes − Desulfotomaculales Incertae Sedis
Firmicutes − Desulfotomaculaceae
Firmicutes − Desulfitobacteriaceae
Firmicutes − Desulfitibacteraceae
Firmicutes − Defluviitaleaceae
Firmicutes − Clostridiaceae
Firmicutes − Christensenellaceae
Firmicutes − Carnobacteriaceae
Firmicutes − Caloramatoraceae
Firmicutes − Caldicoprobacteraceae
Firmicutes − Caldalkalibacillaceae
Firmicutes − Brevibacillaceae
Firmicutes − Bacillaceae
Firmicutes − Aneurinibacillaceae
Firmicutes − Anaerovoracaceae
Firmicutes − Alicyclobacillaceae
Firmicutes − Aerococcaceae
Firmicutes − Acidaminococcaceae
Firmicutes − Acholeplasmataceae
Firmicutes − [Eubacterium] coprostanoligenes group
GS : IG
GS : PC
IG : PC
GS : VC
IG : VC
PC : VC
Sample type comparisons
Adjusted p-values
P ≤ 0.0001(****)
P ≤ 0.001(***)
P ≤ 0.01(**)
P ≤ 0.05 (*)
P > 0.05 (ns)
Sample Comparison
Ø A > Ø B
Ø A < Ø B
Sample type
GS: Gut soil (pooled)
IG: Gut soil (individual)
PC: Pre-Compost
VC: Vermicompost

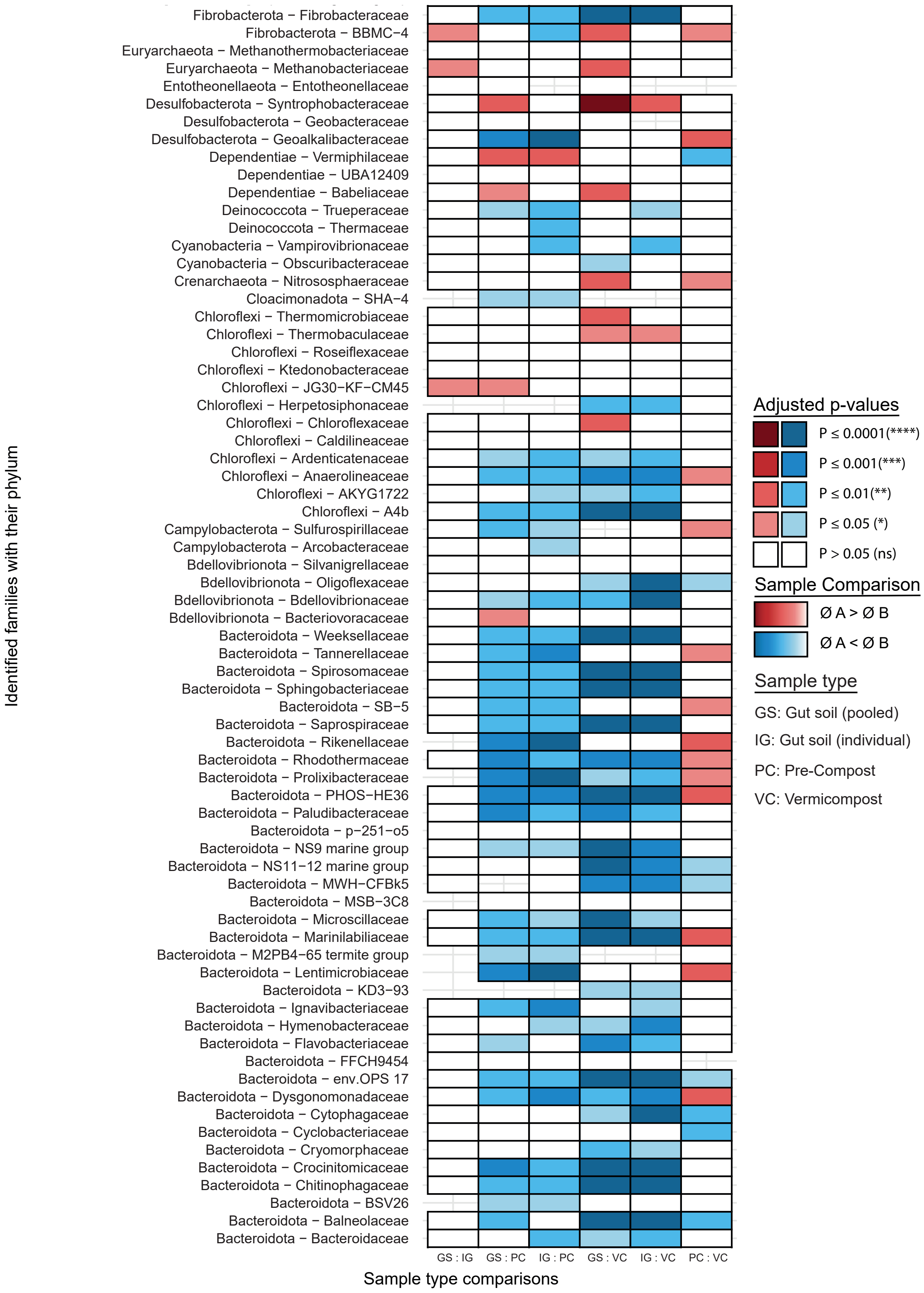

Identified families with their phylum
Fibrobacterota − Fibrobacteraceae
Fibrobacterota − BBMC−4
Euryarchaeota − Methanothermobacteriaceae
Euryarchaeota − Methanobacteriaceae
Entotheonellaeota − Entotheonellaceae
Desulfobacterota − Syntrophobacteraceae
Desulfobacterota − Geobacteraceae
Desulfobacterota − Geoalkalibacteraceae
Dependentiae − Vermiphilaceae
Dependentiae − UBA12409
Dependentiae − Babeliaceae
Deinococcota − Trueperaceae
Deinococcota − Thermaceae
Cyanobacteria − Vampirovibrionaceae
Cyanobacteria − Obscuribacteraceae
Crenarchaeota − Nitrososphaeraceae
Cloacimonadota − SHA−4
Chloroflexi − Thermomicrobiaceae
Chloroflexi − Thermobaculaceae
Chloroflexi − Roseiflexaceae
Chloroflexi − Ktedonobacteraceae
Chloroflexi − JG30−KF−CM45
Chloroflexi − Herpetosiphonaceae
Chloroflexi − Chloroflexaceae
Chloroflexi − Caldilineaceae
Chloroflexi − Ardenticatenaceae
Chloroflexi − Anaerolineaceae
Chloroflexi − AKYG1722
Chloroflexi − A4b
Campylobacterota − Sulfurospirillaceae
Campylobacterota − Arcobacteraceae
Bdellovibrionota − Silvanigrellaceae
Bdellovibrionota − Oligoflexaceae
Bdellovibrionota − Bdellovibrionaceae
Bdellovibrionota − Bacteriovoracaceae
Bacteroidota − Weeksellaceae
Bacteroidota − Tannerellaceae
Bacteroidota − Spirosomaceae
Bacteroidota − Sphingobacteriaceae
Bacteroidota − SB−5
Bacteroidota − Saprospiraceae
Bacteroidota − Rikenellaceae
Bacteroidota − Rhodothermaceae
Bacteroidota − Prolixibacteraceae
Bacteroidota − PHOS−HE36
Bacteroidota − Paludibacteraceae
Bacteroidota − p−251−o5
Bacteroidota − NS9 marine group
Bacteroidota − NS11−12 marine group
Bacteroidota − MWH−CFBk5
Bacteroidota − MSB−3C8
Bacteroidota − Microscillaceae
Bacteroidota − Marinilabiliaceae
Bacteroidota − M2PB4−65 termite group
Bacteroidota − Lentimicrobiaceae
Bacteroidota − KD3−93
Bacteroidota − Ignavibacteriaceae
Bacteroidota − Hymenobacteraceae
Bacteroidota − Flavobacteriaceae
Bacteroidota − FFCH9454
Bacteroidota − env.OPS 17
Bacteroidota − Dysgonomonadaceae
Bacteroidota − Cytophagaceae
Bacteroidota − Cyclobacteriaceae
Bacteroidota − Cryomorphaceae
Bacteroidota − Crocinitomicaceae
Bacteroidota − Chitinophagaceae
Bacteroidota − BSV26
Bacteroidota − Balneolaceae
Bacteroidota − Bacteroidaceae
GS : IG
GS : PC
IG : PC
GS : VC
IG : VC
PC : VC
Sample type comparisons
Adjusted p-values
P ≤ 0.0001(****)
P ≤ 0.001(***)
P ≤ 0.01(**)
P ≤ 0.05 (*)
P > 0.05 (ns)
Sample Comparison
Ø A > Ø B
Ø A < Ø B
Sample type
GS: Gut soil (pooled)
IG: Gut soil (individual)
PC: Pre-Compost
VC: Vermicompost

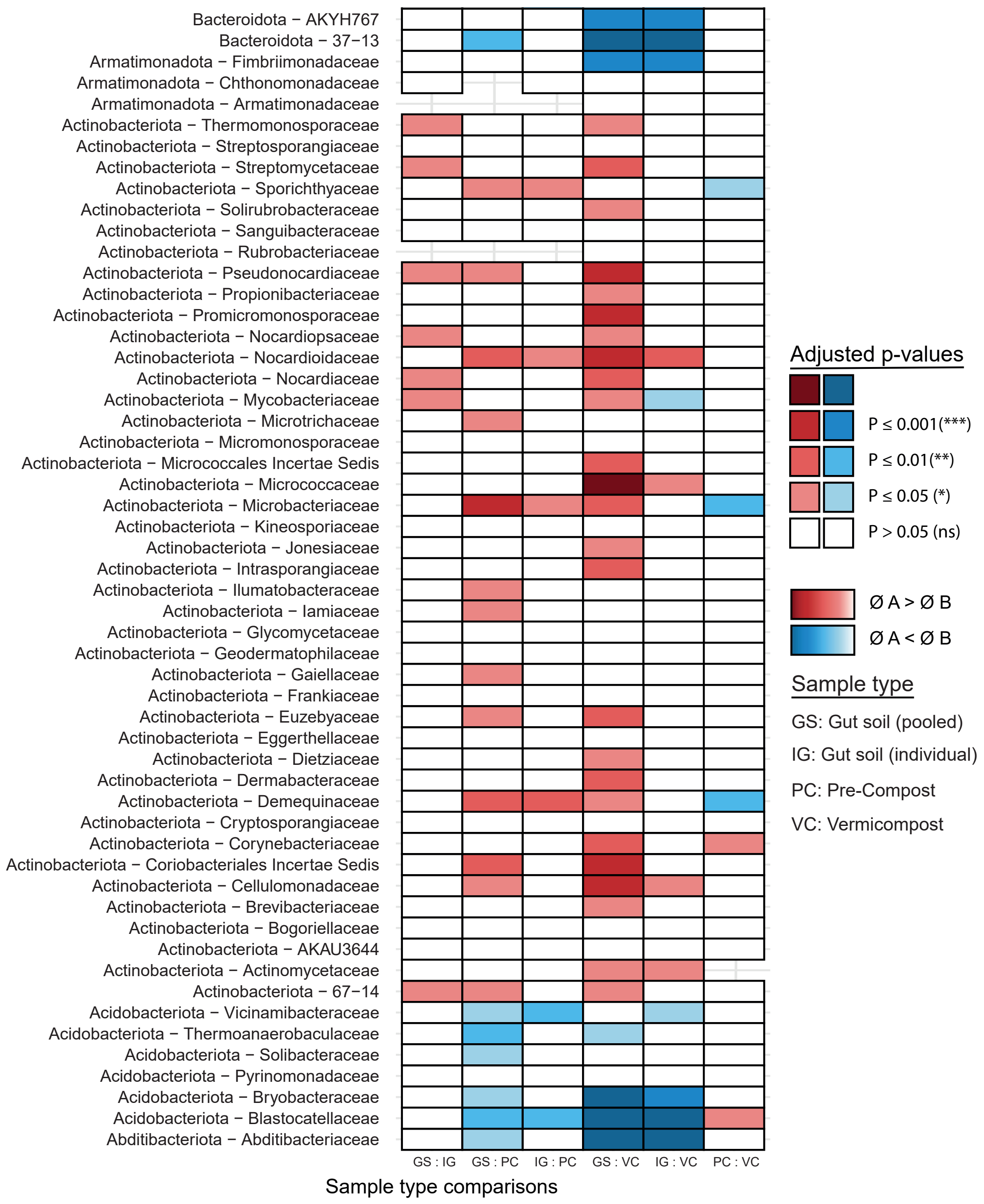


**Figure S2.** Pair wise comparisons of each sample type (GS), individual gut soil (IG), Pre-compost (PC) and Vermicompost (VC) for each family identified by Wilcoxon test with post Benjamini-Hochberg showing adjusted p-values for each comparison (p > 0.05 (ns), p ≤ 0.05 (*), p ≤ 0.01 (**), p ≤ 0.001 (***), p ≤ 0.0001 (****)). Blue color scale indicates that the mean of sample A is lower that in sample B. Red color scale indicates that the mean of sample A is greater than the mean of sample B.

**Figure S3.** Rarefied taxonomic relative abundance plots with notable PUFA producers outlined. (**A**) Prokaryote 16S rRNA rarefied to 3k. (**B**) Eukaryote 18S rRNA with Oligochaetes removed and rarefied to 1k.

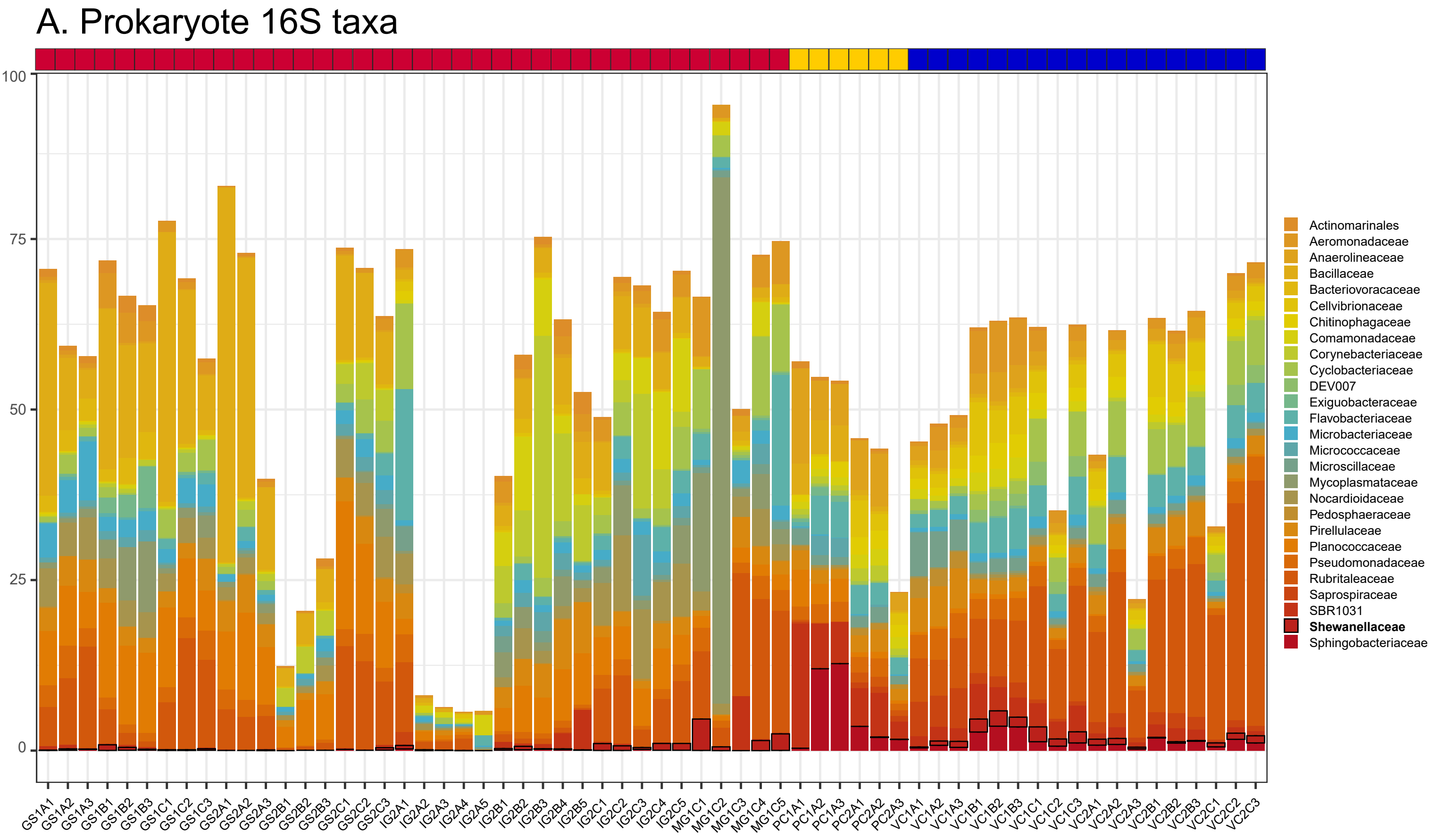


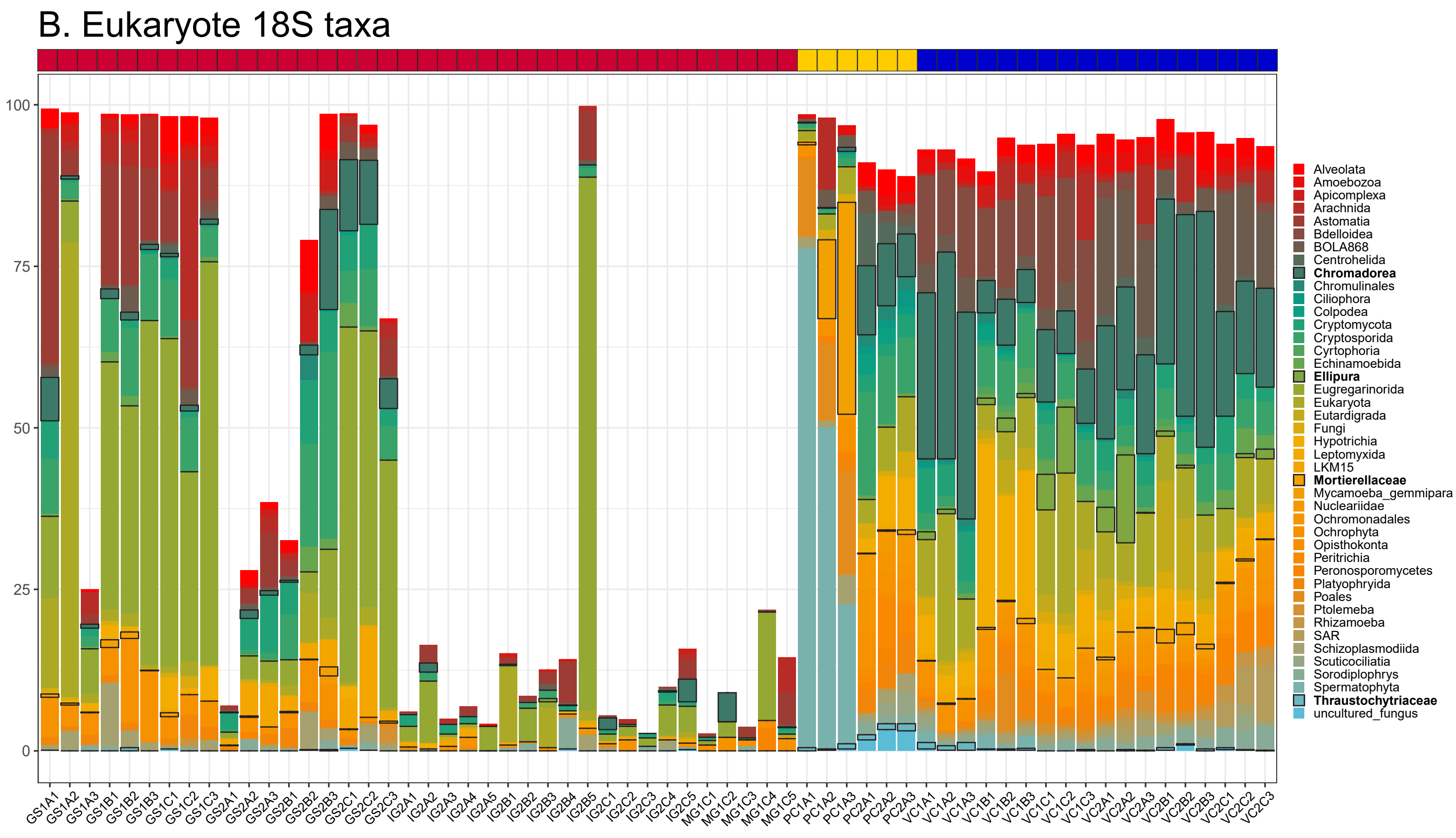

**Figure S4.** Phylogenetic distribution of PfaA-KS primer matches

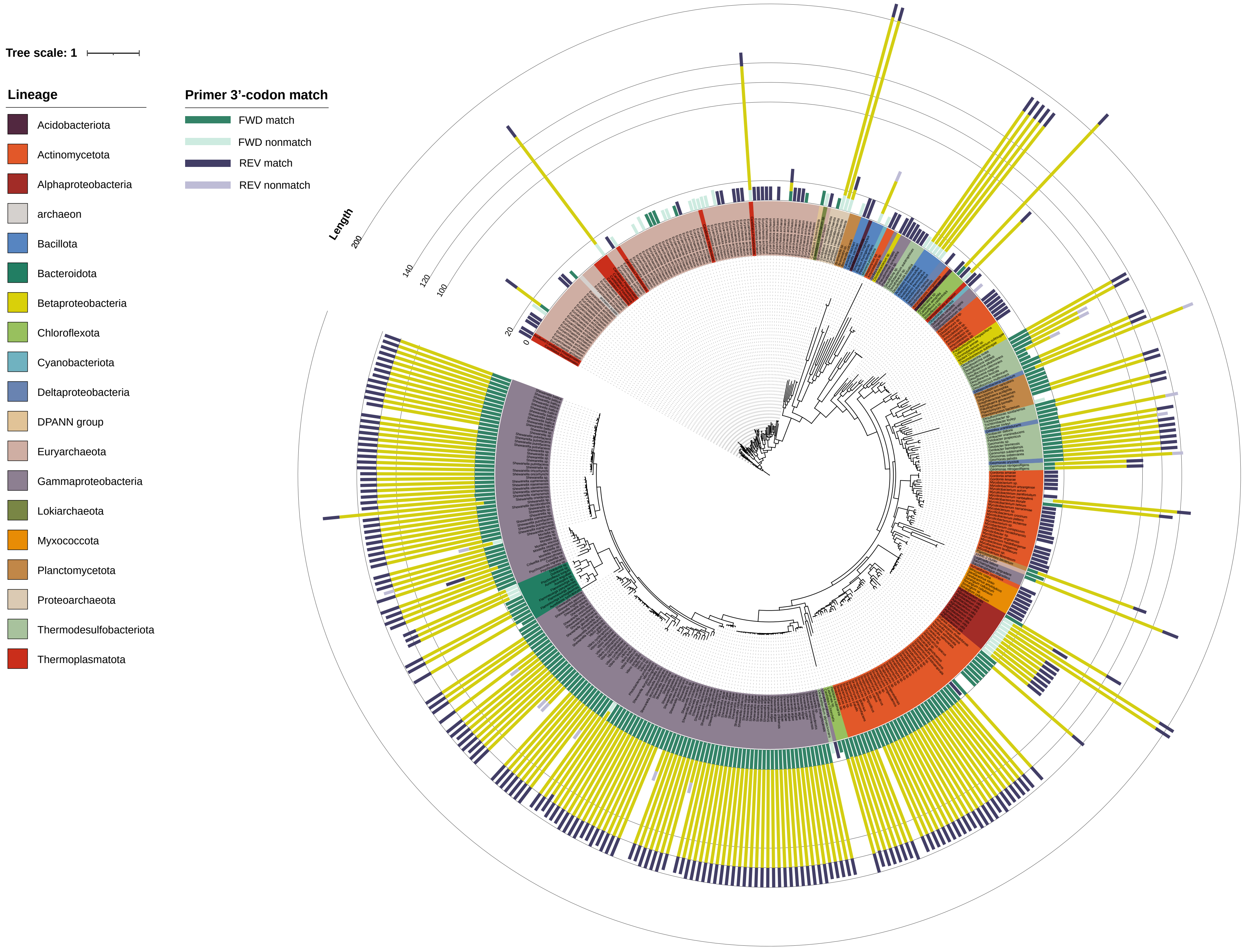

**Figure S5.** Associations between total PfaA-KS ASVs and metadata

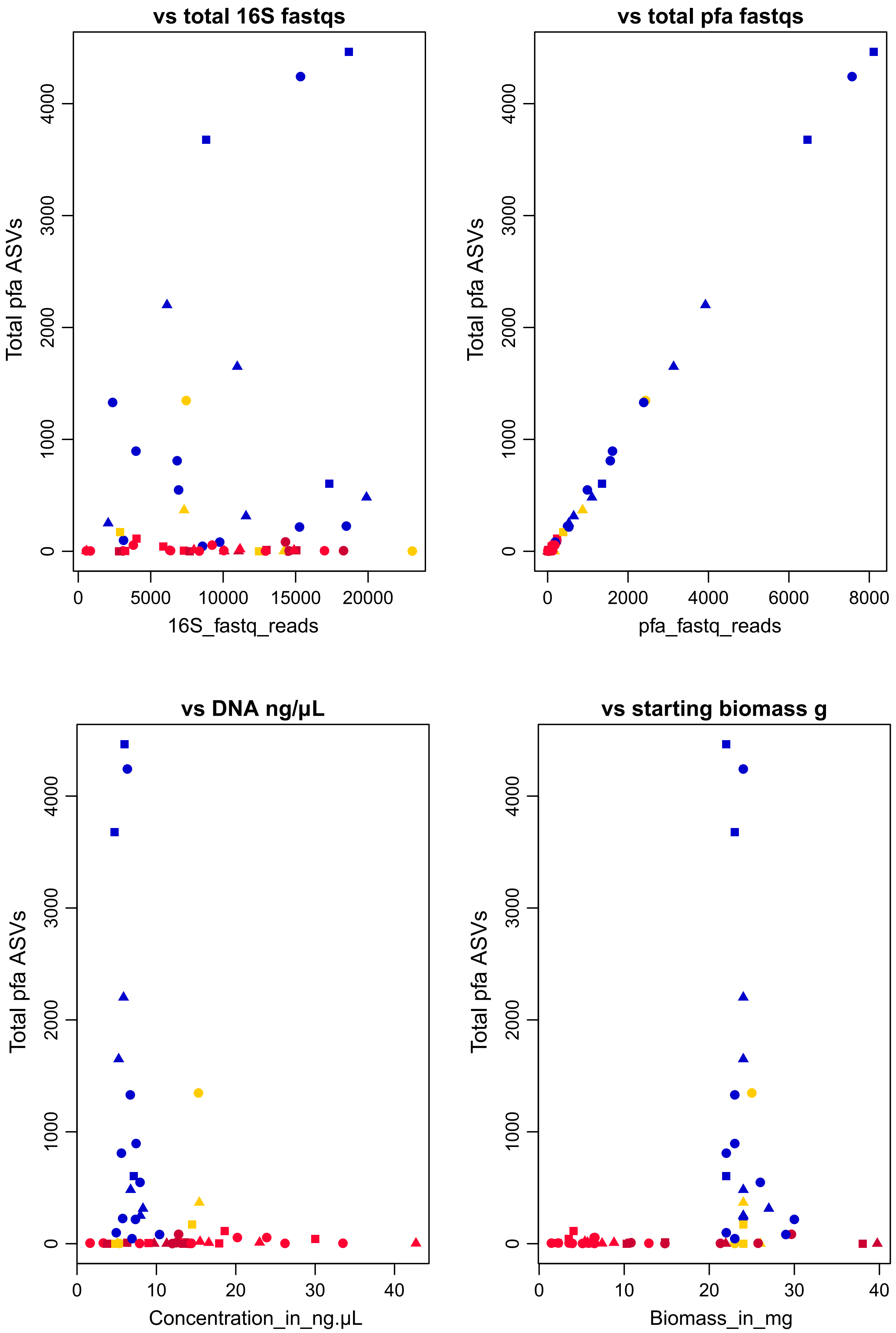

**Figure S6**. Abundance of PUFA taxa among 16S rRNA data

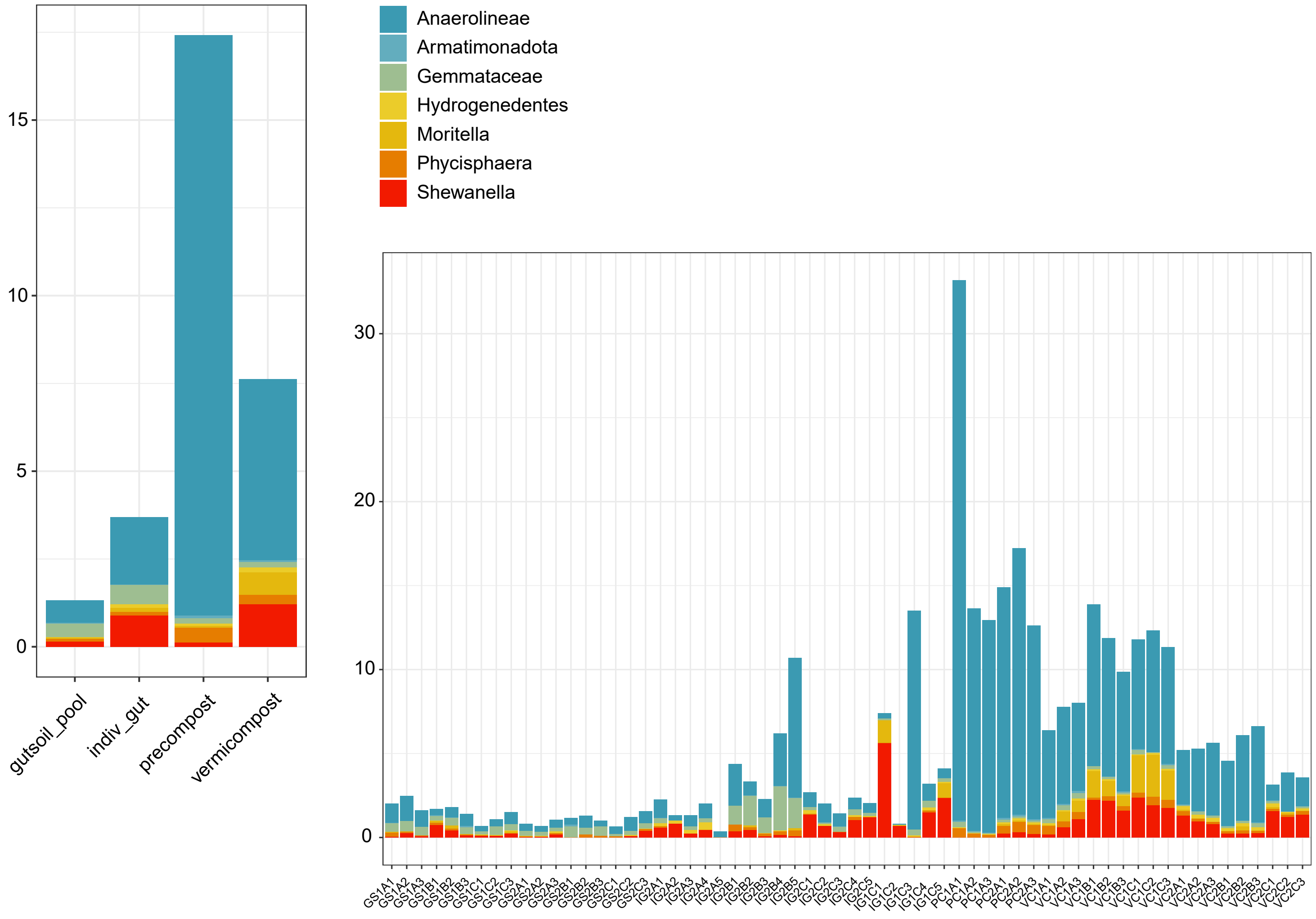

**Figure S7.** Mantel's test dotplot of **pfa** and **prok.pfa** datasets showing comparison of pairwise distances

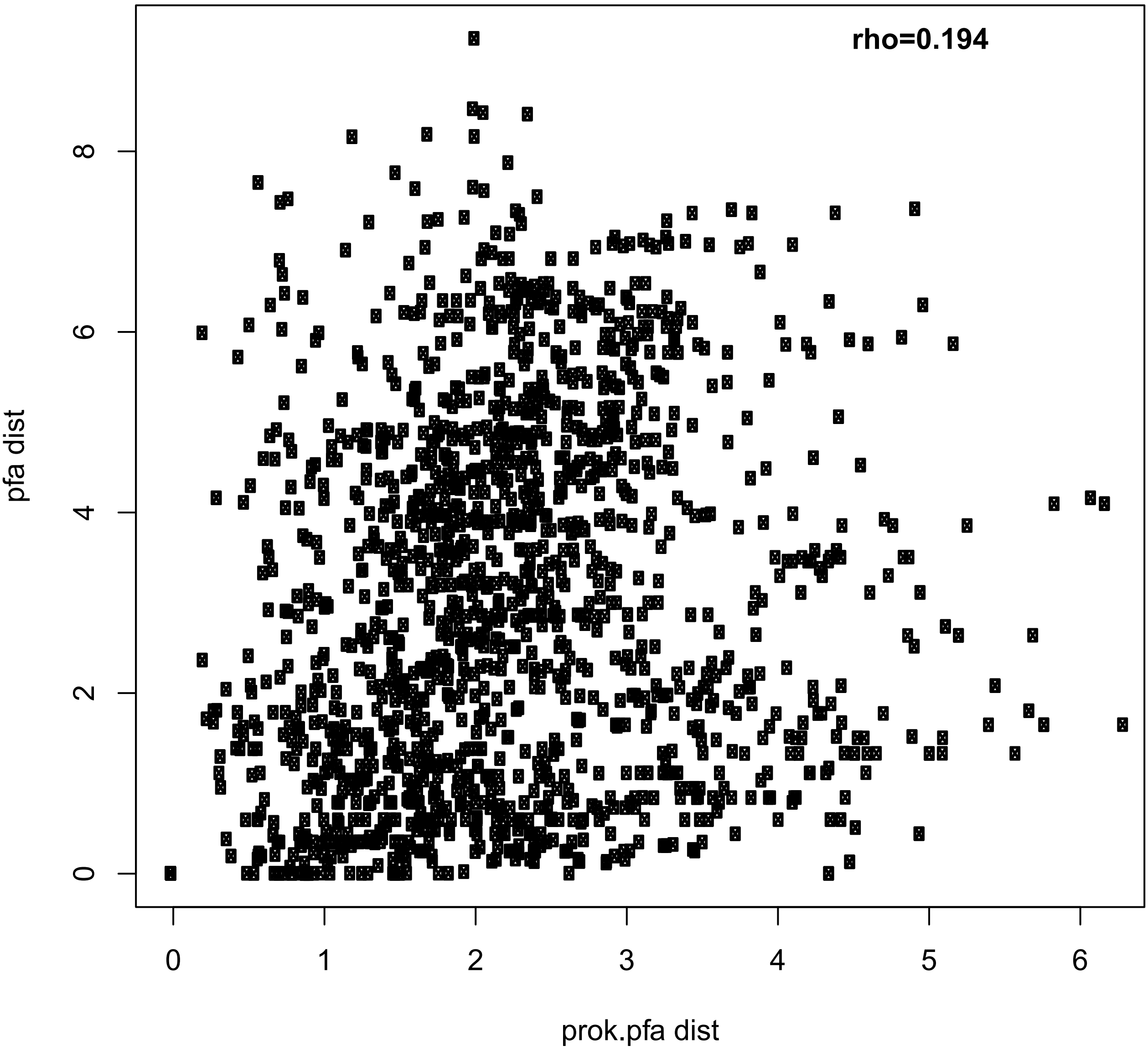

**Figure S7.** Linear associations between taxonomic abundance for pfa and prok.pfa datasets

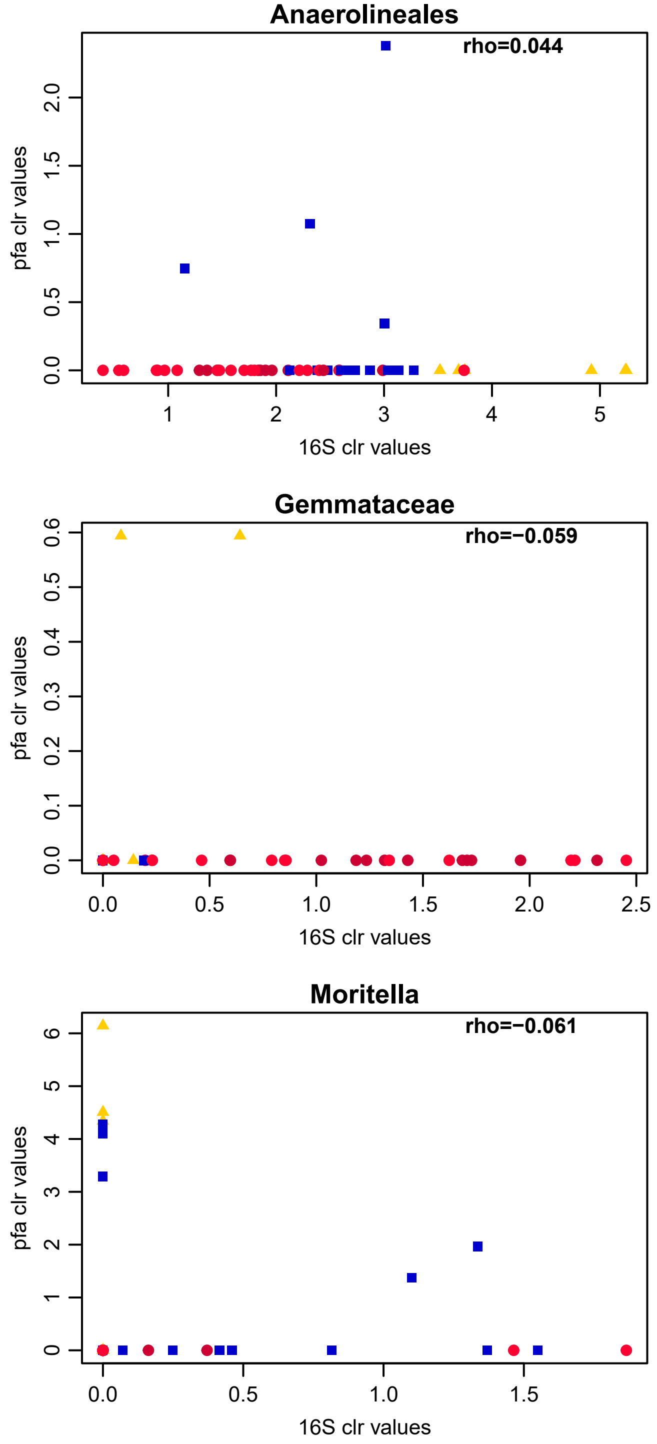


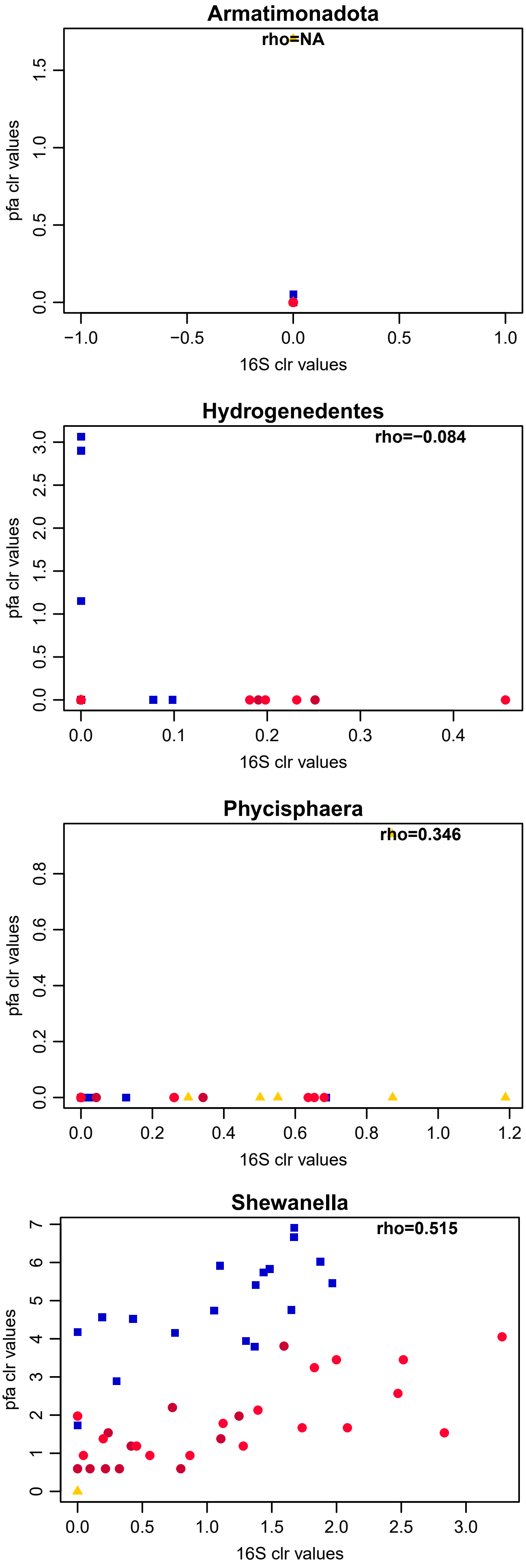

**Figure S9.** Distributions for pfa taxa in both datasets

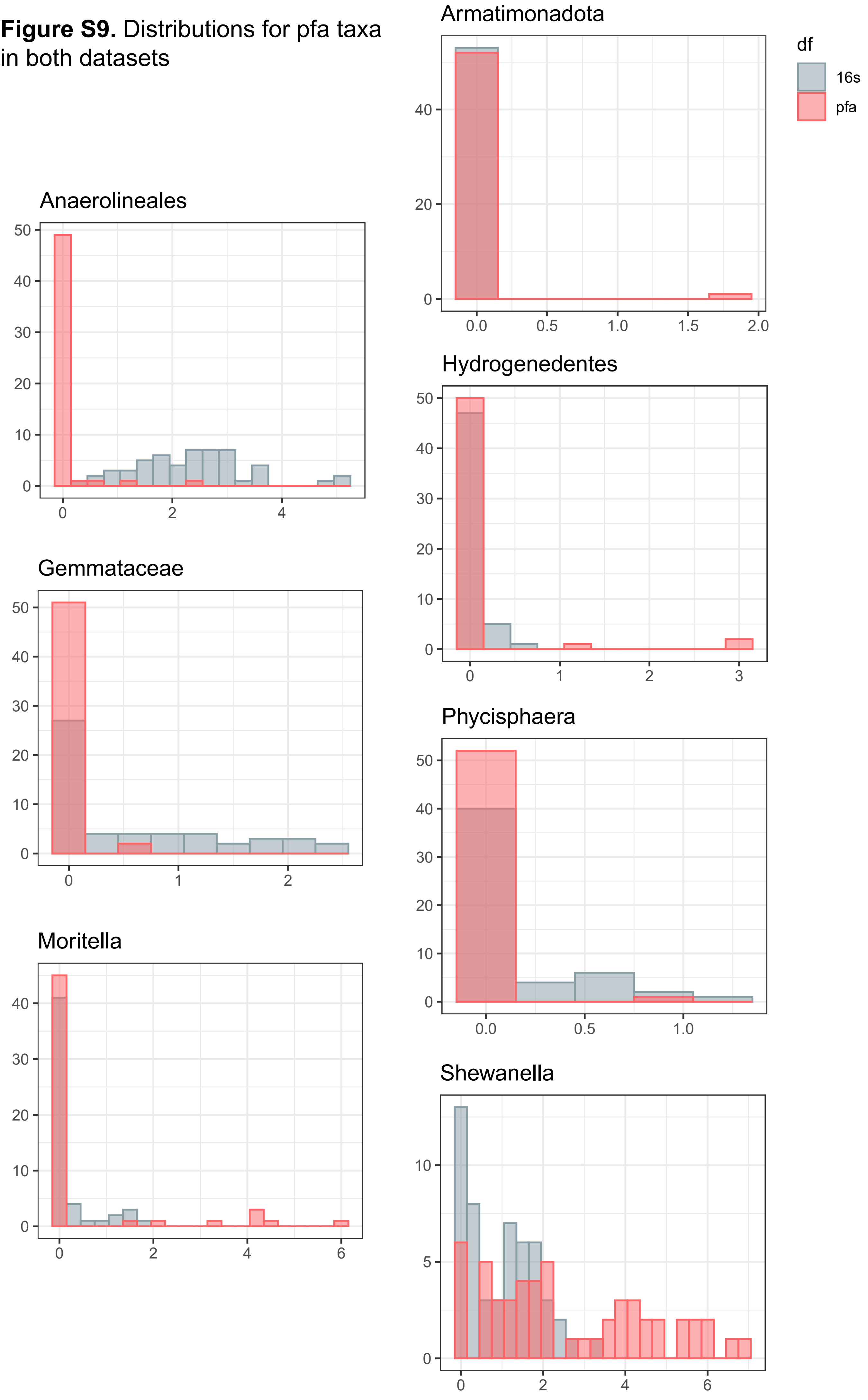

**Figure S10.** Effect size pairwise comparisons of the transformed abundances of pfa-taxa drawn from the prokaryote 16S dataset and compared across sample types

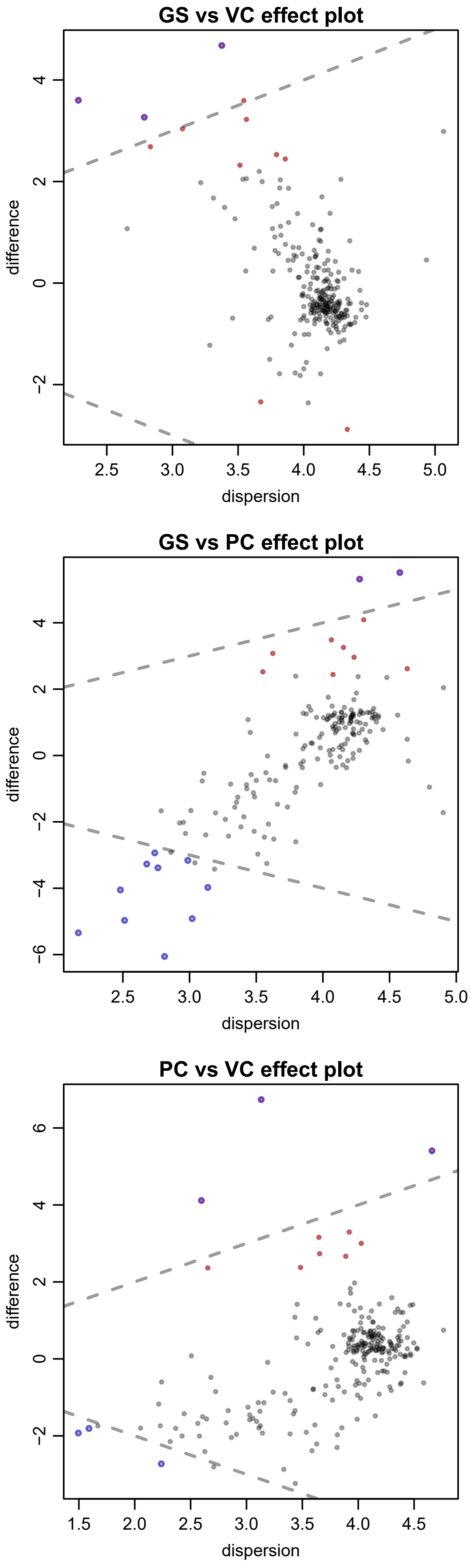
